# Prediction of residue-residue contacts in CASP12 targets from its predicted tertiary structures

**DOI:** 10.1101/192120

**Authors:** Piyush Agrawal, Sandeep Singh, Gandharva Nagpal, Deepti Sethi, Gajendra P.S. Raghava

## Abstract

One of the challenges in the field of structural proteomics is to predict residue-residue contacts in a protein. It is an integral part of CASP competitions due to its importance in the field of structural biology. This manuscript describes RRCPred 2.0 a method participated in CASP12 and predicted residue-residue contact in targets with high precision. In this approach, firstly 150 predicted protein structures were obtained from CASP12 Stage 2 tarball and ranked using clustering-based quality assessment software. Secondly, residue-residue contacts were assigned in top 10 protein structures based on distance between residues. Finally, residue-residue contacts were predicted in target protein based on consensus/average in top 10 predicted structures. This simple approach performs better than most of CASP12 methods in the categories of TBM and TBM/FM. It ranked 1^st^ in following categories; i) TBM domain on list size L/5, ii) TBM/FM domain on list size L/5 and iii) TBM/FM domain on Top 10. These observations indicate that predicted tertiary structure of a protein can be used for predicting residue-residue contacts in protein with high accuracy.

## Introduction

Numbers of methods have been developed in the past for predicting three-dimension (3D) structure of a protein from its primary structure or amino acid sequence. Broadly, these approaches can be classified in three classes; i) template or homology based approach, ii) threading based approach and iii) *ab initio* method^1^. In recent years, *ab initio* method is gaining importance in the field of structure prediction since traditional method of template based structure prediction is limited by the number of structural templates in the Protein Data Bank (RCSB-PDB)^2^. However, recent studies showed residue-residue contact map guided structure prediction, as an important intermediate step in *ab initio* structure prediction of a protein^3,4^. Apart from structure prediction, residue-residue contact predictions has also shown promising application in the field of drug designing^5^, model ranking, selection and its evaluation^6,7^. Although the importance of residue-residue contact prediction in modeling tertiary structure of a protein was introduced decades ago^8,9^, the thorough implementation of the idea came into practice when several groups across the world have shown how residues in contact can be predicted with improved and reasonable accuracy^10,11^. One of the main reasons of development in this field is introduction of a separate section of residue-residue contact prediction in CASP (Critical Assessment of Techniques for Protein Structure Prediction). This section was introduced in the CASP2^12^ but it was fully standardized during CASP6-CASP9^13-16^. Definition of residue-residue contact has not been changed since previous CASP rounds. Two residues are said to be in contact when distance between their Cβ atoms (Cα in case of glycine) is shorter than 8.0 Å. These contacts are also classified in three different categories: short, medium and long range contacts based on separation along the sequence. Short range contacts are those contacts where residues are separated by 6-11 residues, Medium range contacts are those where residues are separated by 12-23 residues whereas Long range contacts are those where residues are separated by minimum 24 residues or more than that^17^. Among all the contacts, ‘long range contact’ prediction is of much interest and more valuable for structure prediction^18^. These contacts are evaluated using precision i.e. number of contacts correctly predicted out of all the predicted contacts^19^

Current existing methods on residue-residue contact prediction can be classified into two categories. First is sequence based method, where no information other than sequence of a protein is present. Contacts in the protein are predicted utilizing information which can be derived directly from its sequence. Number of methods have been developed utilizing various machine-learning techniques such as support vector machines^20,21^, hidden markov models^22^, neural networks^23,24^, deep network and boosting techniques^25^, evolutionary information^26,27^, covariation signal information^28^, etc. Second class is template/structure-based methods where structure is used for extracting contact information. In this method, firstly structure of protein is modeled using best available template and then contacts are predicted from the modeled structure^21^.

As reported in literature, prediction of contacts depends very much on the quality of the template structure^21^. Therefore, keeping this in mind, in the present study, we have developed a simple approach to predict residue-residue contact from high quality tertiary structure predicted by various methods participated in structure prediction category of CASP12. We ranked all predicted structure and used top 10 structures for predicting residue-residue contact using consensus or average based approach.

## Results and Discussion

A large number of groups participated in CASP12 residue-residue contact prediction category, where they submitted their predicted models to CASP12 site (http://predictioncenter.org/casp12/doc/CASP12_Abstracts.pdf). In this paper, we analyzed and discussed performance of models manually submitted by our group “raghavagps”. Our prediction models were derived from Stage 2 tarballs released by CASP12 sites. These tarballs comprises models submitted by online servers to CASP12 site (See Materials & Methods). In this CASP12, we have analyzed the results present at the site (http://predictioncenter.org/casp12/rrc_results.cgi). Here, our result was evaluated on 66 targets out of total 94 targets for ‘long’ and ‘long+medium’ range contacts by the CASP team. Out of our 66 targets, 38 targets were present in Free Modeling (FM) category, 12 in Template Based Modeling (TBM) category and 16 in Template Based Modeling/Free Modeling (TBM/FM) category.

### Prediction of ‘long’ range Contacts

Firstly, we analyzed the prediction performance of all groups in predicting ‘long’ range contacts on targets of TBM domain category (Table 1). The performance of models were extracted from CASP12 sites using probability >0 on list size L/5 (Supplementary Fig. S1). As shown in Table 1, method RRCPred 2.0 (raghavagps) got average precision value of 85.028%, which is better than other methods in TBM category. This method RRCPred 2.0 (raghavagps) was followed by “naive” and “FLOUDAS_SERVER”, with 2^nd^ and 3^rd^ position respectively. Similarly, performance was extracted from CASP site for Top10 contacts for all groups on targets of TBM category. On Top 10 contacts method “naive” got average precision value of 86.486%, which is better than other methods (Table 1). As shown in Table 1, method “naïve” was followed by “RaptorX-Contact” and RRCPred 2.0 (raghavagps) with 2^nd^ and 3^rd^ rank respectively (Supplementary Fig. S2).

**Table 1:** The performance or precision of different methods for ‘long’ range contacts on CASP12 TBM targets.

| Methods/groups | L/5* |  | Top10 |  |
| --- | --- | --- | --- | --- |
|  | Precision (%) | Rank | Precision (%) | Rank |
| RRCPred 2.0 (raghavagps) | 85.028 | 1 | 83.333 | 3 |
| Naïve | 80.247 | 2 | 86.486 | 1 |
| FLOUDAS_SERVER | 77.321 | 3 | 77.941 | 6 |
| Deepfold-Contact | 76.801 | 4 | 82.432 | 4 |
| RaptorX-Contact | 76.745 | 5 | 83.784 | 2 |
| Distill | 75.051 | 6 | 77.568 | 7 |
| iFold_1 | 72.105 | 7 | 77.500 | 8 |
| Yang-Server | 72.088 | 8 | 79.444 | 5 |
| MetaPSICOV | 71.582 | 9 | 76.571 | 9 |
| PconsC31 | 69.672 | 10 | 75.405 | 10 |

\* where L is the sequence length of the protein.

In addition to TBM targets, we also extracted the performance of different methods on targets falling in TBM/FM category using probability value >0. The precision of predicting ‘long’ range contact of different methods for L/5 is shown in Table 2. In this category (TBM/FM) for L/5, method RRCPred 2.0 (raghavagps) achieved precision 72.244% with recall 16.466%, which is better than other methods. This method is followed by “iFold_1” and “MetaPSICOV” with precision 58.117% and 53.755% respectively (Supplementary Fig. S3). Similar trend was observed for Top10 contacts where method RRCPred 2.0 (raghavagps) achieved precision 73.125%, followed by “iFold_1” and “PconsC31” which got precision 62.353% and 61.579% respectively (Supplementary Fig. S4). It is important to note that performance of all methods/groups decreased substantially for TBM/FM targets in comparison to TBM targets.

**Table 2.** The performance or precision of different methods for ‘long’ range contacts on CASP12 TBM/FM targets.

| Methods | L/5* |  | Top10 |  |
| --- | --- | --- | --- | --- |
|  | Precision (%) | Rank | Precision (%) | Rank |
| RRCPred 2.0 (raghavagps) | 72.244 | 1 | 73.125 | 1 |
| iFold_1 | 58.117 | 2 | 62.353 | 2 |
| MetaPSICOV | 53.755 | 3 | 59.474 | 4 |
| PconsC31 | 52.342 | 4 | 61.579 | 3 |
| Naive | 51.988 | 5 | 54.211 | 7 |
| Deepfold-Contact | 51.026 | 6 | 53.684 | 8 |
| myprotein-me | 47.892 | 7 | 55.000 | 6 |
| RaptorX-Contact | 46.579 | 8 | 47.368 | 12 |
| RBO-Epsilon | 46.286 | 9 | 57.368 | 5 |
| Pcons-net | 45.721 | 10 | 50.526 | 10 |

\* where L is the sequence length of the protein.

Most challenging targets in CASP competition are targets of FM category. We extracted the performance of all methods from CASP site for FM targets. As shown in Table 3, method “RaptorX-Contact” with average precision of 47.095% in L/5 list size outperformed other methods. The performance of all methods was very poor for FM category targets. The method “RaptorX-Contact” was followed by “iFold_1” and “MetaPSICOV” who scored 2^nd^ and 3^rd^ rank respectively. Unfortunately, our method RRCPred 2.0 (raghavagps) performed very poor on FM targets and achieved precision value of 32.883% and ranked 17^th^ (Supplementary Fig. S5). We also analysed the performance of different methods for Top10 contacts on FM targets and observed similar trends (Supplementary Fig. S6). This means method RRCPred 2.0 (raghavagps) is not suitable for FM category targets.

**Table 3.** The performance or precision of different methods for ‘long’ range contacts on CASP12 FM targets.

| Methods | L/5<sup>*</sup> |  | Top10 |  |
| --- | --- | --- | --- | --- |
|  | Precision (%) | Rank | Precision (%) | Rank |
| RaptorX-Contact | 47.095 | 1 | 54.211 | 1 |
| iFold_1 | 46.864 | 2 | 50.270 | 2 |
| Deepfold-Contact | 45.549 | 3 | 48.889 | 4 |
| MetaPSICOV | 44.007 | 4 | 47.368 | 6 |
| MULTICOM-CLUSTER | 41.691 | 5 | 48.947 | 3 |
| MULTICOM-CUNSTRUCT | 41.637 | 6 | 48.421 | 5 |
| Naive | 41.388 | 7 | 45.556 | 8 |
| Pcons-net | 41.013 | 8 | 46.579 | 7 |
| IGBteam | 38.784 | 9 | 43.421 | 9 |
| Zhang_Contact | 37.801 | 10 | 40.833 | 13 |
| RRCPred 2.0 (raghavagps) | 32.883 | 17 | 37.632 | 18 |

<sup>\*</sup> where L is the sequence length of the protein.

When we selected TBM and TBM/FM options collectively, we observed that our method ranked 1^st^ again in both the list size L/5 as well and Top10 with precision of 77.723% and 77.500% respectively (Supplementary Fig. S7 and S8 respectively). In this section, method “naïve” secured 2^nd^ position. When domain TBM/FM and FM selected collectively, our method performed little low in comparison to other methods and ranked 6^th^ with average precision of 44.546% in L/5 list size (Supplementary Fig. S9) and 8^th^ with average precision of 48.148% in Top10 list size (Supplementary Fig. S10). Method “iFold_1” secured 1^st^ position in this section with average precision of >50.00% in both the list size category. In the case of TBM and FM domain, our method didn’t performed so well and showed average precision of 45.398% and 48.600% in L/5 and Top10 list size respectively (Supplementary Fig. S11 and S12 respectively). Method “RaptorX-Conatct” outperformed other methods in this section. When all the three domains were selected at a given time and performance was evaluated, our methods showed descent performance with average precision of 51.906% in L/5 list size (Supplementary Fig. S13) and 54.545% in Top10 list size (Supplementary Fig. S14). In this section method “Deepfold-Contact” ranked 1^st^ in L/5 list size category with average precision of 59.249% whereas method “RaptorX-Conatct” ranked 1^st^ in L/5 list size category with average precision of 64.468%.

### Prediction of ‘long+medium’ range contacts

We extracted the performance of methods on ‘long+medium’ range contacts from CASP12 site for TBM targets using probability >0 on list size L/5. As shown in Table 4, our method RRCPred 2.0 (raghavagps) achieved average precision of 89.101 and stood 1^st^ in the list (Supplementary Fig. S15), it was followed by “naive” and “RaptorX-Contact” at 2^nd^ and 3^rd^ rank respectively. In case of Top10 list size “naive” got 1^st^ rank with average precision 91.892% and was followed by RRCPred 2.0 (raghavagps) and “RaptorX-Contact” with average precision of 89.167% and 88.378% at 2^nd^ and 3^rd^ rank respectively (Supplementary Fig. S16).

**Table 4.**
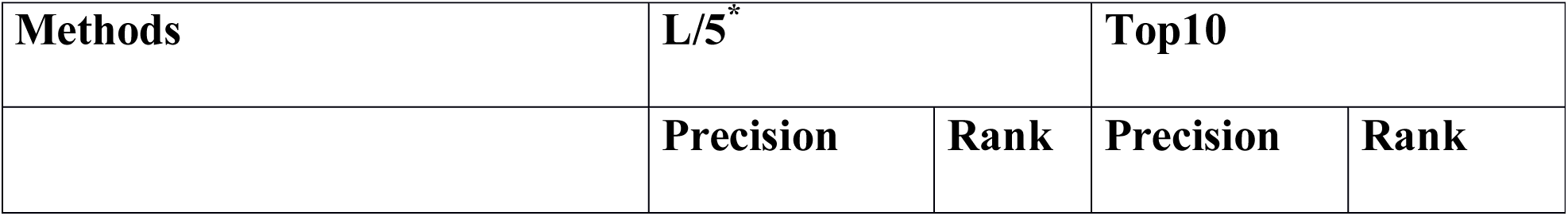
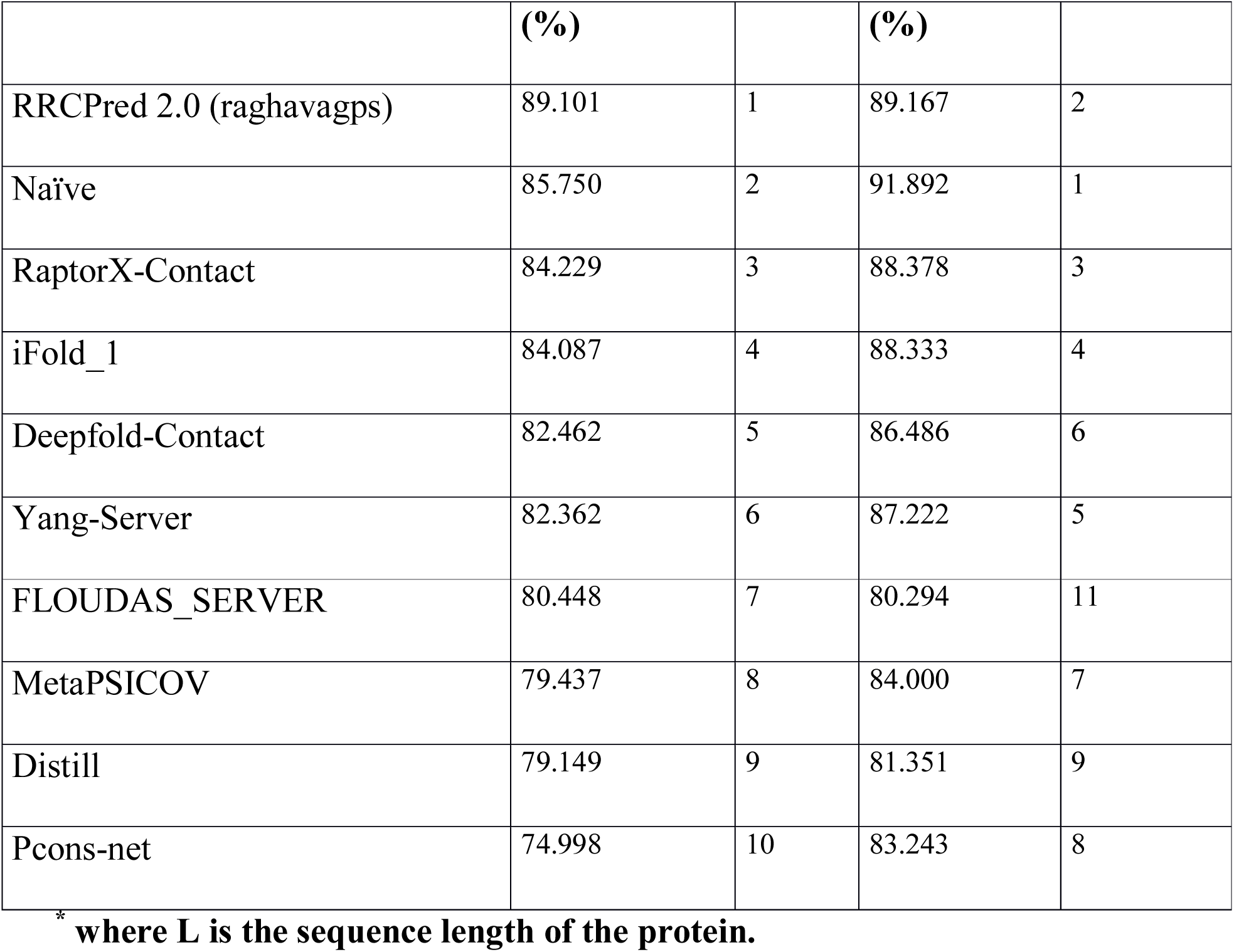
The performance or precision of different methods for ‘long+medium’ range contacts on CASP12 TBM targets.

In addition to TBM targets, we also analyzed the performance of different methods on TBM/FM targets for list size L/5 and Top10 and probability >0 (Table 5). In this category (TBM/FM), RRCPred 2.0 (raghavagps) obtained average precision of 78.358% for L/5 list size which is better than other methods (Supplementary Fig. S17). “MetaPSICOV” was present at 2^nd^ position and “RaptorX-Contact” at 3^rd^ position for L/5 list size. Similar trend was observed for Top10 list size where RRCPred 2.0 (raghavagps) achieved precision of 79.375% (Supplementary Fig. S18) and performed better than other methods.

**Table 5.**
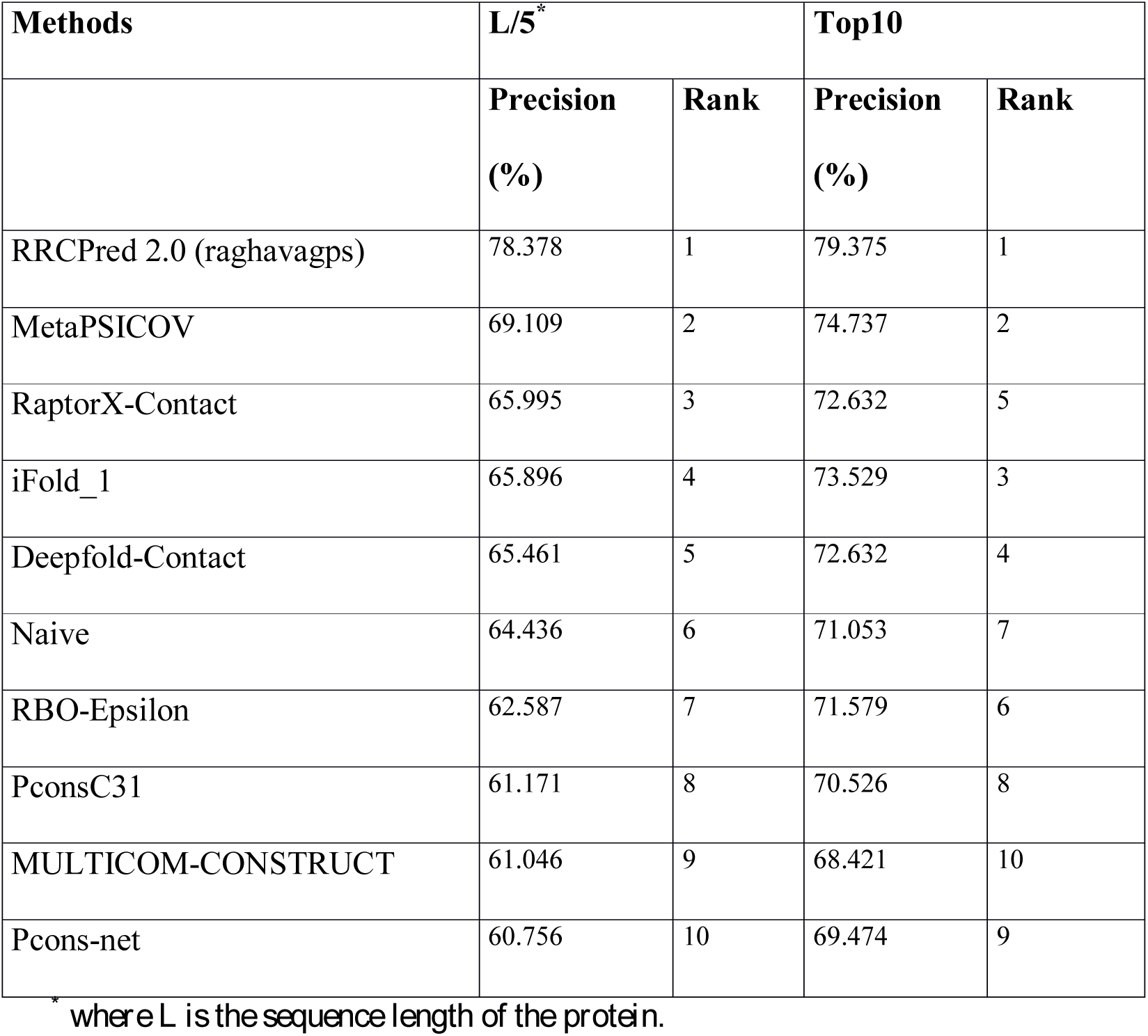
The performance or precision of different methods for ‘long+medium’ range contacts on CASP12 TBM/FM targets.

In case of FM domain targets as shown in Table 6, our method didn’t performed well in both list size L/5 (Supplementary Fig. S19) and Top10 list size (Supplementary Fig. S20) categories. Here, method “RaptorX-Contact” outperformed other methods with average precision of 55.831% in L/5 list size and 63.158% in Top10 list size.

**Table 6.**
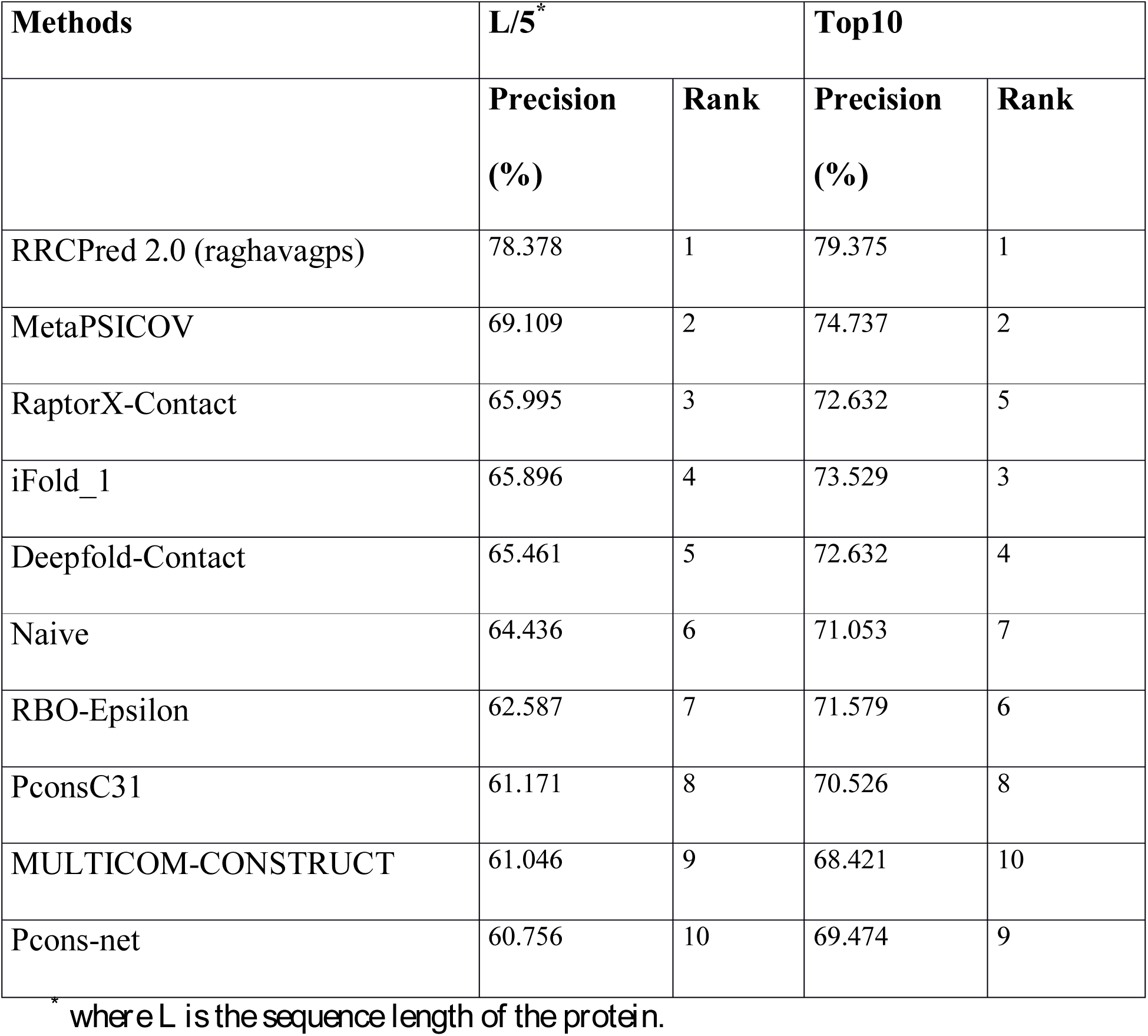
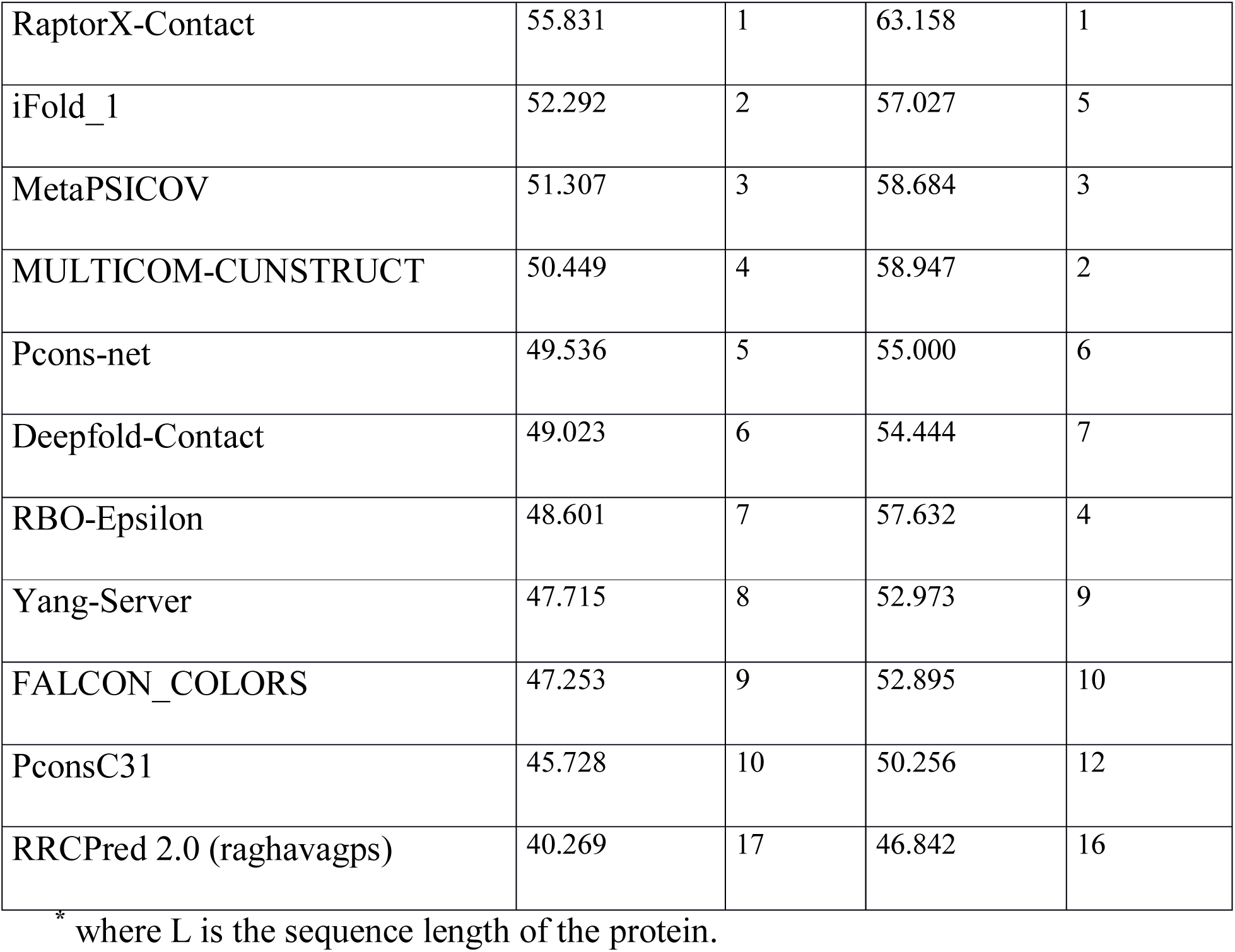
The performance or precision of different methods for ‘long+medium’ range contacts on CASP12 FM targets.

In case of TBM and TBM/FM domain selection, our method showed highest average precision of 82.974% when list size L/5 was selected (Supplementary Fig. S21), followed by method “naive” and “RaptorX-Contact” at 2^nd^ and 3^rd^ ppsition respectively. In list size Top10, method “naive” ranked 1^st^ with average precision of 84.821% and our method RRCPred 2.0 (raghavagps) ranked second with average precision value of 83.571% (Supplementary Fig. S22). When the domain selection TBM and TBM/FM was changed to TBM/FM and FM, our method perform little poor in comparison to other methods and ranked 8^th^ in L/5 list size (Supplementary Fig. S23) whereas 11^th^ in Top10 list size (Supplementary Fig. S24). Method “RaptorX-Contact” outperformed other methods in both list size. In the case of TBM and FM domain, our method showed average precision value of 51.989% and 57.000% in L/5 (Supplementary Fig. S25) and Top10 list size (Supplementary Fig. S26). “RaptorX-Contact” was found to be at the top with average precision of 69.840% in L/5 and 75.600% in Top10 list size. When all the three domains were selected, our method showed reasonable performance of average precision value of 58.386% (Supplementary Fig. S27) in L/5 list size and 62.424% in Top10 list size (Supplementary Fig. S28).

### Target wise analysis

In addition to average performance of methods, we analyzed the performance of our method target wise. The performance for each target using different options were extracted from CASP12 site using probability >0. As shown in Table 7, for a large number of targets our method got 100% precision. In case of ‘long’ range contacts for L/5, “raghavagps” achieved more than 70% precision for 28 targets that includes 4 targets with 100% precision (Supplementary Table S1). This performance improved for Top10 contacts, where our method predicts 10 targets with 100% precision (Supplementary Table S2). This shows utility of our method in real life. As expected, performance on ‘long+medium’ range was better than ‘long’ range contacts. In case of ‘long+medium’ range contacts for L/5 our method achieved more than 70% for 31 targets with 5 targets showing precision of 100% (Supplementary Table S3). In case of Top10 for ‘long+medium’ range contacts, our method predicts 38 targets with more than 70% precision (Supplementary Table S4). For more detailed information, see (Supplementary Table S5-S8) where performances of all methods have been compiled for L/5 ‘long’ range contacts (Supplementary Table S5), Top10 ‘long’ range contacs (Supplementary Table S6), L/5 ‘long+medium’ range contacts (Supplementary Table S7) and Top10 ‘long+medium’ range contacts (Supplementary Table S8) respectively.

**Table 7:**
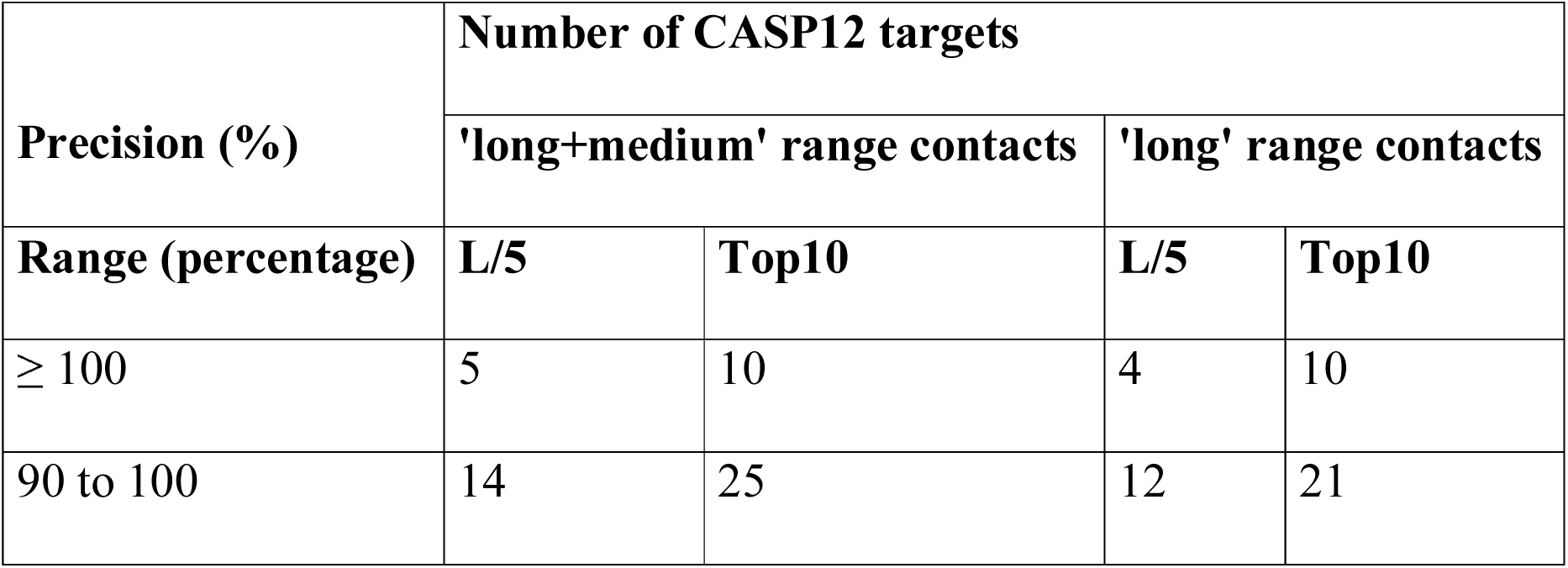
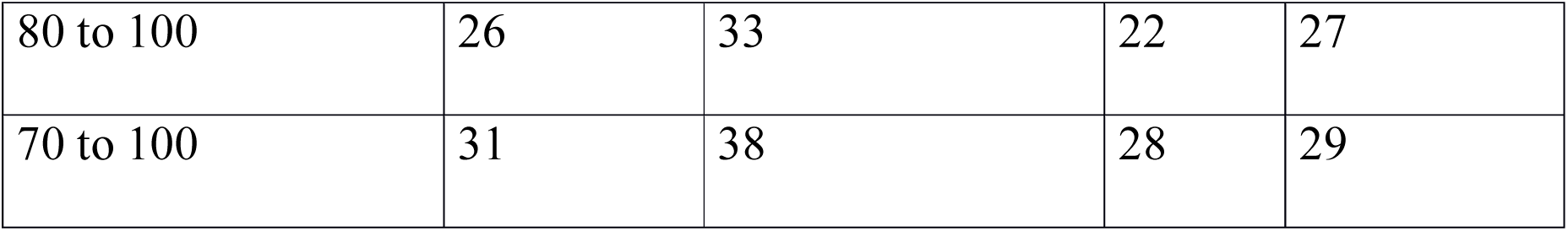
Summary of target-wise performance of method RRCPred 2.0 (raghavagps) on long range and long+medium range contact.

### Evaluation and ranking by CASP12 organizers

Once competition is over, CASP organizers evaluate the performance of methods participated in competition and provides the results in the form of presentation; CASP12 results, analyzed by the organizers is also available from its website (http://predictioncenter.org/casp12/doc/presentations/CASP12_RR-Gaeta.pdf). We found that the organizers have analyzed the results on FM category targets where the common parameters were contacts which was of ‘medium’ and ‘long’ range combined and SumZ (>-2) whereas varying parameters were list size (L/2, L/5 and Full list), probability (>0 and >0.5). Weights taken into consideration were Precision, F1, F1prob., and AUC1. Method “RaptorX-Contact” was found to be present at top position in most of the evaluations and hence becoming one of the promising methods for FM targets contact prediction. We found that our method RRCPred 2.0 (raghavagps) also performed well and was present among the top 10 methods in most of the evaluations. The result suggests that our method performed comparably on FM targets when compared to other top methods and can be improved further. Unfortunately, the performance of TBM and TBM/FM targets were not evaluated and provided by the organizers in the presentation. This is the reason our method was not considered in successful predictors.

## Conclusion

In last two decades, number of methods has been developed for predicting residue-residue contact. Despite tremendous efforts made by scientific community, the performance of these methods is far from satisfactory^29^. In particular, prediction of ‘long’ range contact is a tedious job. Due to its importance, it is highly desirable to develop highly accurate methods for predicting residue-residue contacts in a protein from amino acids sequence. This is the reason that residue-residue contact prediction is part of CASP competition. One of the major goals of CASP is to assess the quality of tertiary structure predicted by different methods and groups. Overall aim is to improve the performance of tertiary structure prediction methods. This raises a question whether one can use the predicted tertiary structures for predicting residue-residue contact in a protein. In order to address this question, our group ‘raghavagps’ used 150 predicted tertiary structures released by CASP12 for each target from Stage 2 tarballs. We ranked these structures using quality assessment software QASproCL then top 10 structures were used to assign/predict residue-residue contact in a target (see Materials & Methods). These predicted residue-residue contacts were submitted to CASP12 in manual submission category. In order to avoid any biasness, we extracted performance of models for all groups from CASP12 web site. This paper presents performance of method RRCPred 2.0 (raghavagps) and its comparison with performance of other methods. As shown in Results section in comparison to other models, this method performs better than other methods or in top 3 methods for both categories TBM and TBM/FM. This means predicted tertiary structure can be used to predict residue-residue contacts with high precision, particularly for easy targets (TBM or TBM/FM). The performance of RRCPred 2.0 (raghavagps) was poor for FM targets; still it is in list of top methods. We used QASproCL for ranking predicted structures, which was not the best method for ranking predicted structure as evaluated by CASP12. Thus it is possible to improve the performance of this method if we used best quality assessment method for ranking predicted structure.

## Methodology

### Dataset and Algorithm

We extracted Stage 2 tarball for each target from CASP12 web site, where each tarball consists of best 150 server models (predicted structures). These models/structures were predicted by servers which participated in TS category in CASP12. These tarballs were given as input in the quality assessment (QA) category in CASP12 competitions and were open to everyone for downloading from their web site. QASproCL, an in-house developed clustering based quality assessment software, was used for computing quality of each model for a given CASP12 target [**http://predictioncenter.org/casp12/doc/CASP12_Abstracts.pdf**]. These models were ranked based on their QASproCL model quality score (MQ) where the model getting the highest quality score was ranked first.. In order to compute residue-residue contact in a protein target, we used top/best 10 models. Following equation 1 was used for computing residue-residue contact score between two residues in a protein.

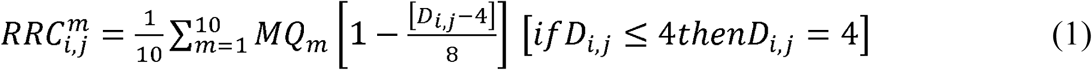

Where m is the mth model, i and j are two residues forming a pair.

where *RRC*_*i,j*_ is residue-residue contact score, *MQ*_*m*_ is model quality of *m*^*th*^ model as predicted by QASproCL and *D*_*i*_ is distance between two residues in Angstrom (Å).

Schematic representation of complete algorithm applied in method “raghavagps” is presented in the form of flowchart (Figure 1).

**Figure 1.**
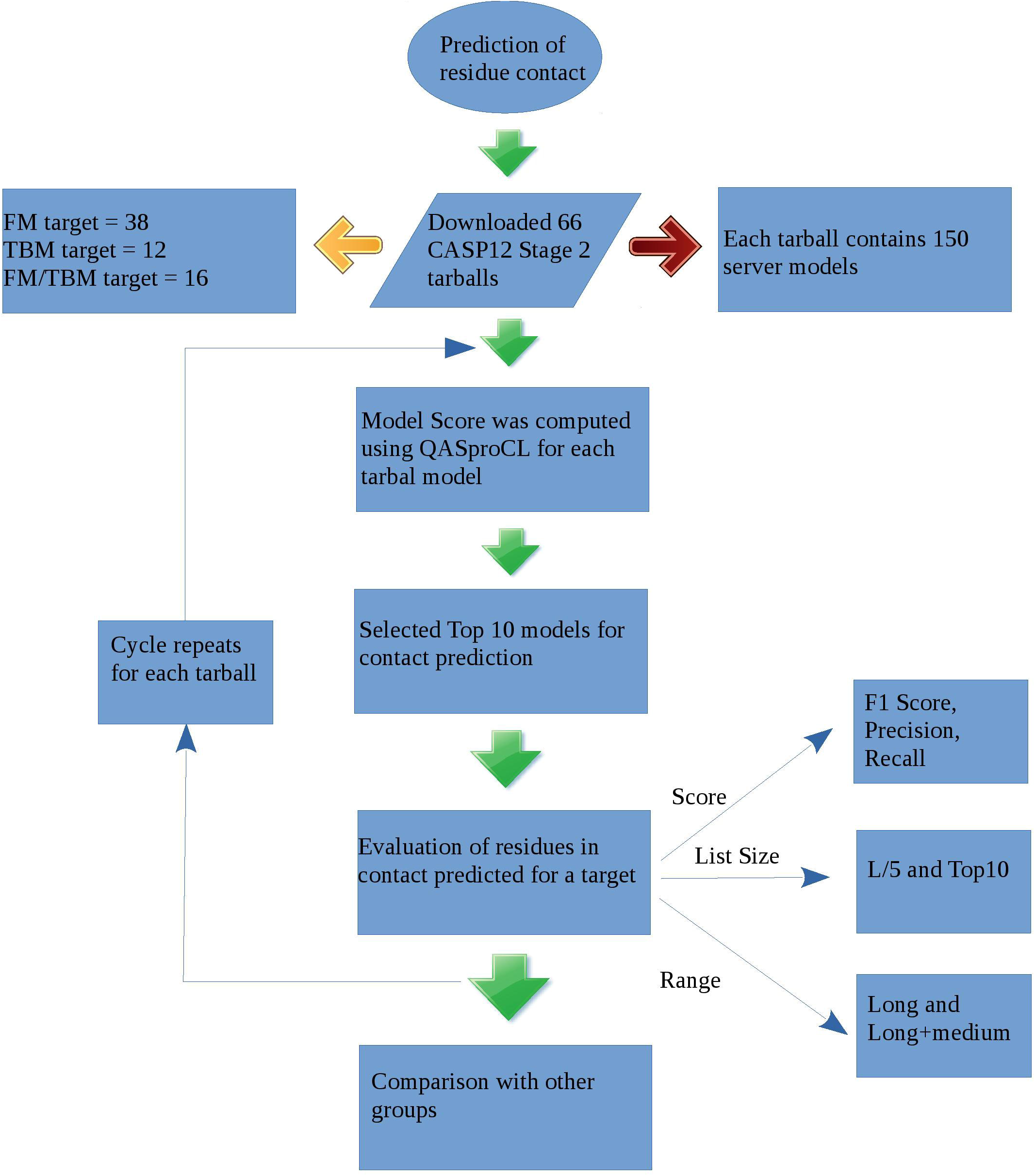
Schematic represntation of the methodolgy followed for predicting residue-residue contact in the method RRCPred 2.0 (raghavagps).

*Insert Figure 1 here*

### Parameters for Evaluation

The performance of prediction models was measured using different parameters (e.g. F1 score, precision, recall), which has been used in earlier CASP competition’s for evaluating group performance^17^. These parameters can be computed using following equation 2-4.

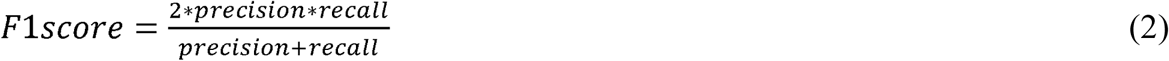

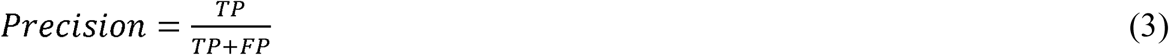

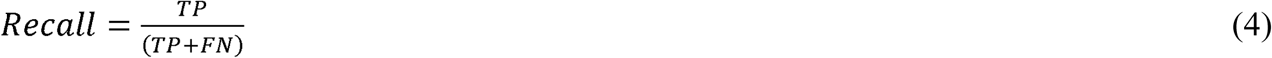

Where *TP* is true positives or correctly predicted contacts, *FP* is false positive or non-contacts being predicted as contacts, *TN* is true negative or correctly predicted non-contacts and FN is false negative or contacts being predicted as non-contacts.

### Criteria for Evaluation

In the present CASP12 competition, ‘long’ and ‘long+medium’ range contacts were analyzed since ‘long’ range contacts are hard to predict. In order to evaluate performance of all contact predictors, number of contacts was kept same by trimming the list to N contacts with highest probabilities. The List Size parameter N determines contact lists length and takes different values i.e. L/2, L/5, Top 10 and FL (Full Length). Here L represents the length of protein sequence; Top 10 means first 10 contacts submitted for a given sequence whereas FL represents all predicted contacts, which is different for different predictions. Different types of scores for each target as well as for each group were calculated for evaluating performance.

## Acknowledgement

Authors are thankful to Dr. Chinmayee Choudhury for her kind help during QASproCL algorithm development and funding agencies Department of Science and Technology (DST-INSPIRE), Council of Scientific and Industrial Research (CSIR), J.C.Bose National Fellowship from Department of Science and Technology (DST), for fellowships and financial support.

## AUTHORS’ CONTRIBUTION

S.S., G.N., P.A. and D.S. developed the algorithm QASproCL. P.A. and S.S extracted the data and predicted the contacts. P.A. submitted the results manually to the CASP site. P.A. extracted the results from CASP12 site and anlayzed the whole data with the help of G.P.S.R. P.A. and G.P.S.R. wrote the manuscript. G.P.S.R. conceived the idea and coordinated the project.

## Supplementary Information

**Figure S1.**
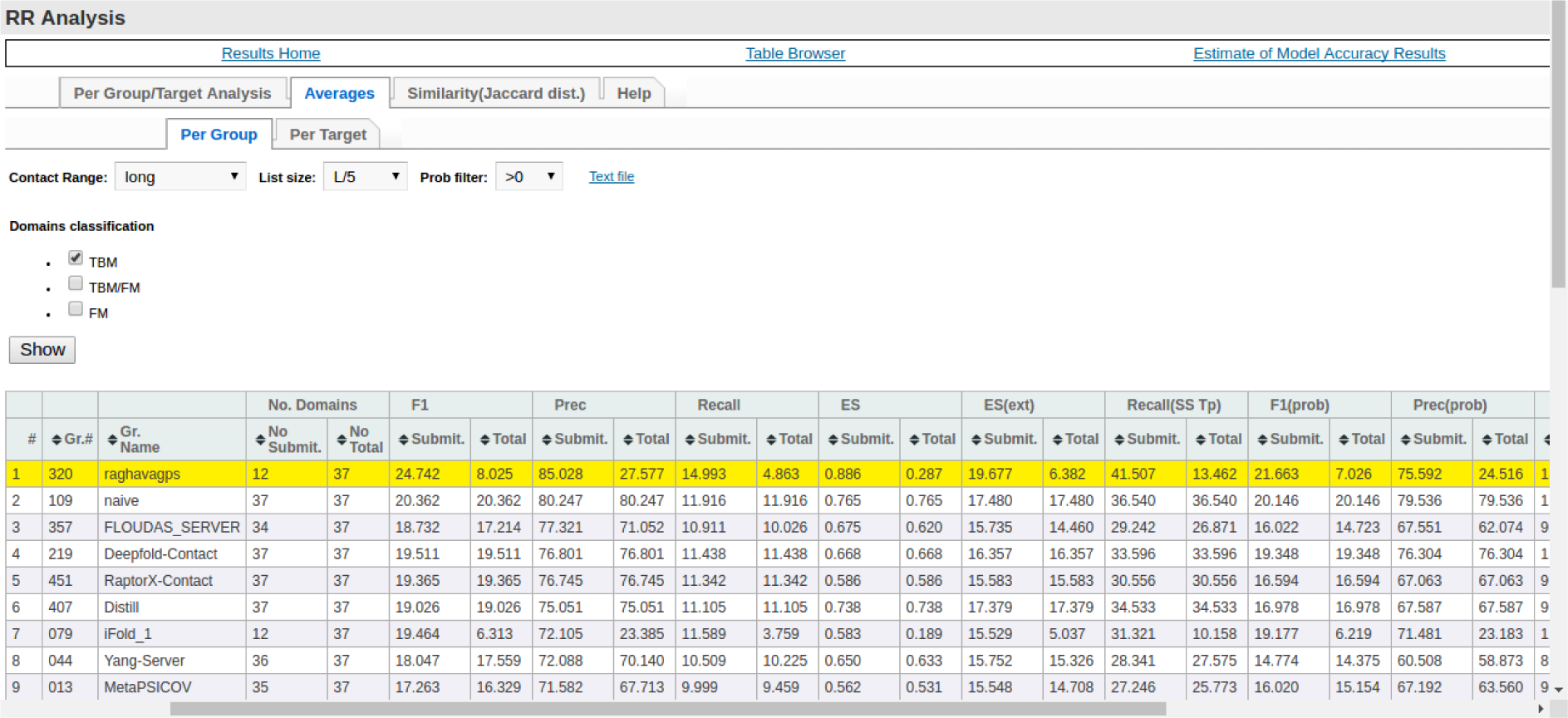
Screenshot of the performance or precision of different methods for L/5 list size and ‘long’ range contacts on CASP12 TBM targets.

**Figure S2.**
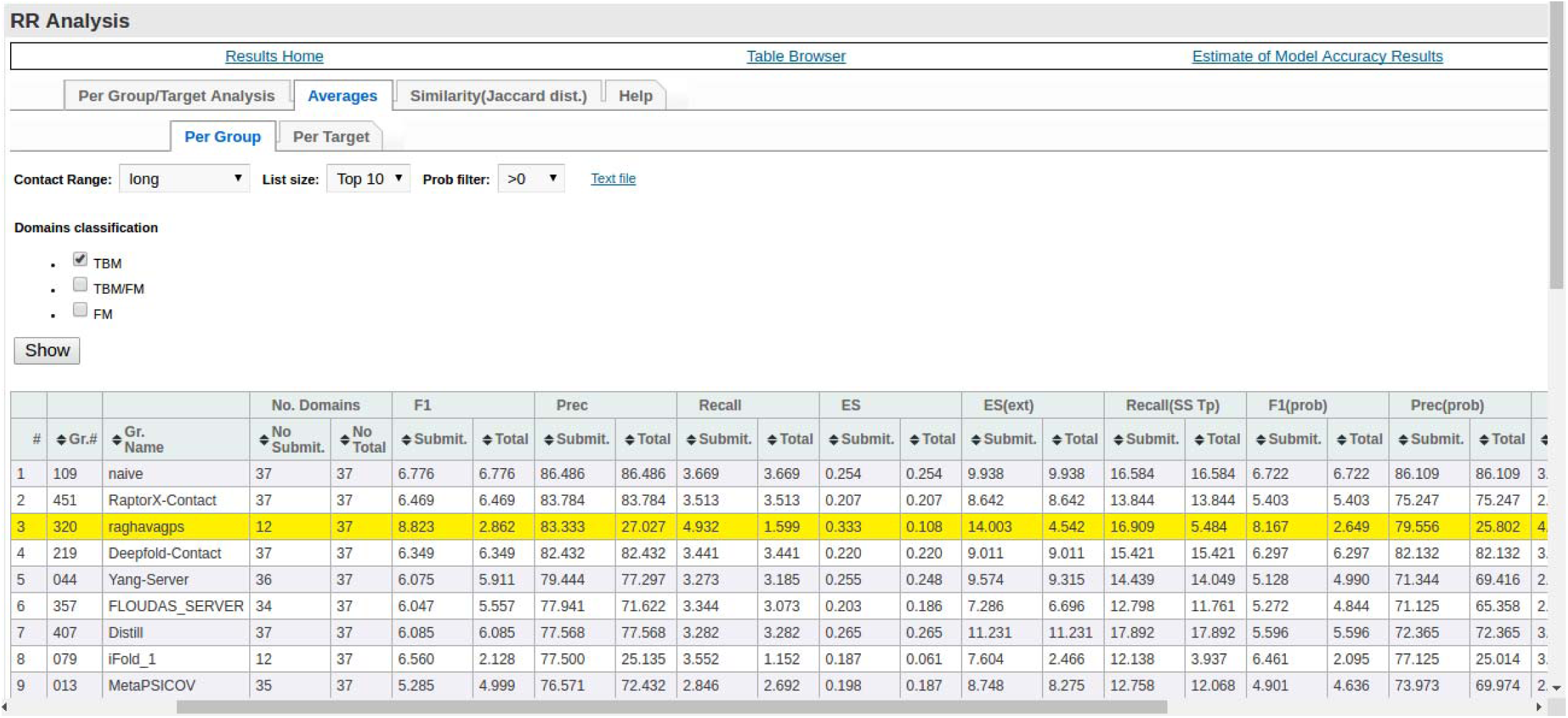
Screenshot of the performance or precision of different methods for Top10 list size and ‘long’ range contacts on CASP12 TBM targets.

**Figure S3.**
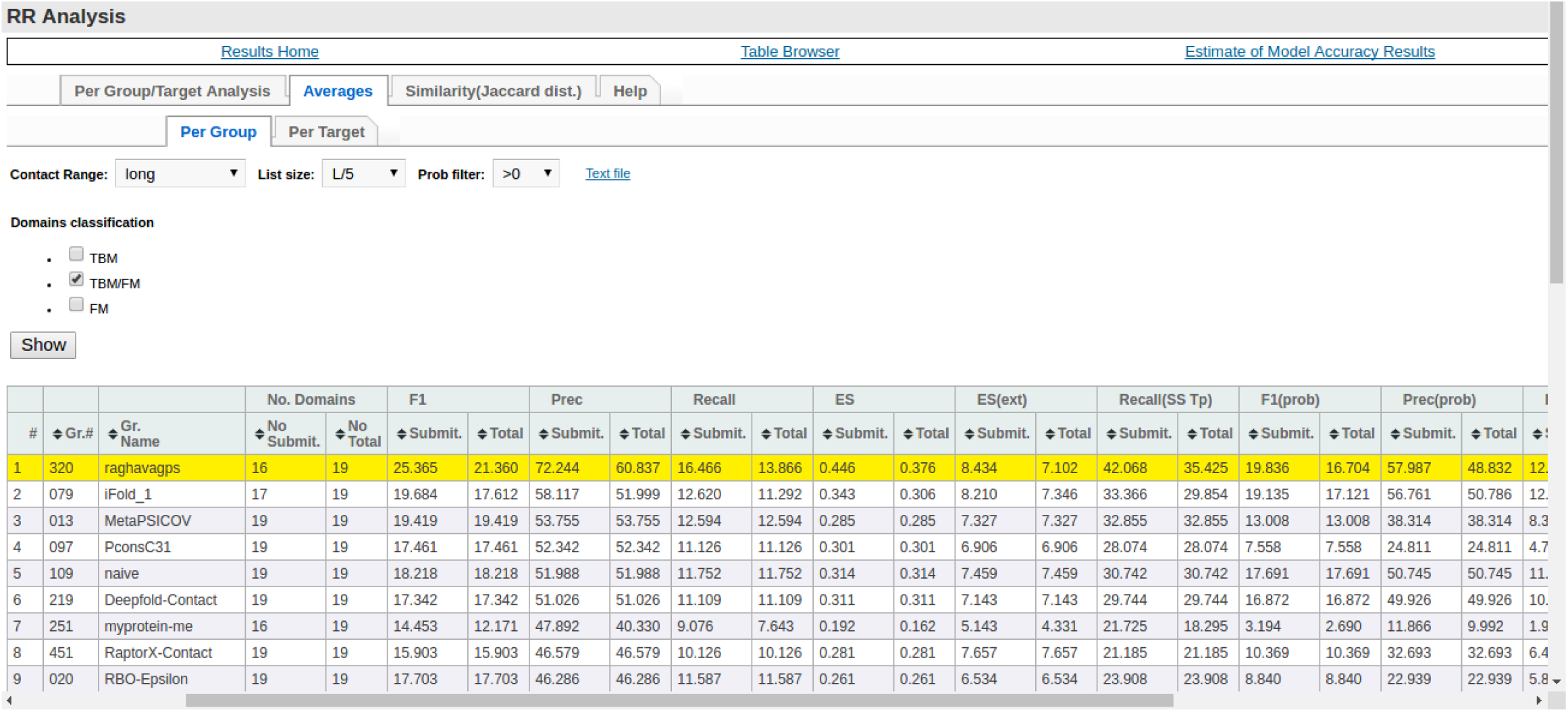
Screenshot of the performance or precision of different methods for L/5 list size and ‘long’ range contacts on CASP12 TBM/FM targets.

**Figure S4.**
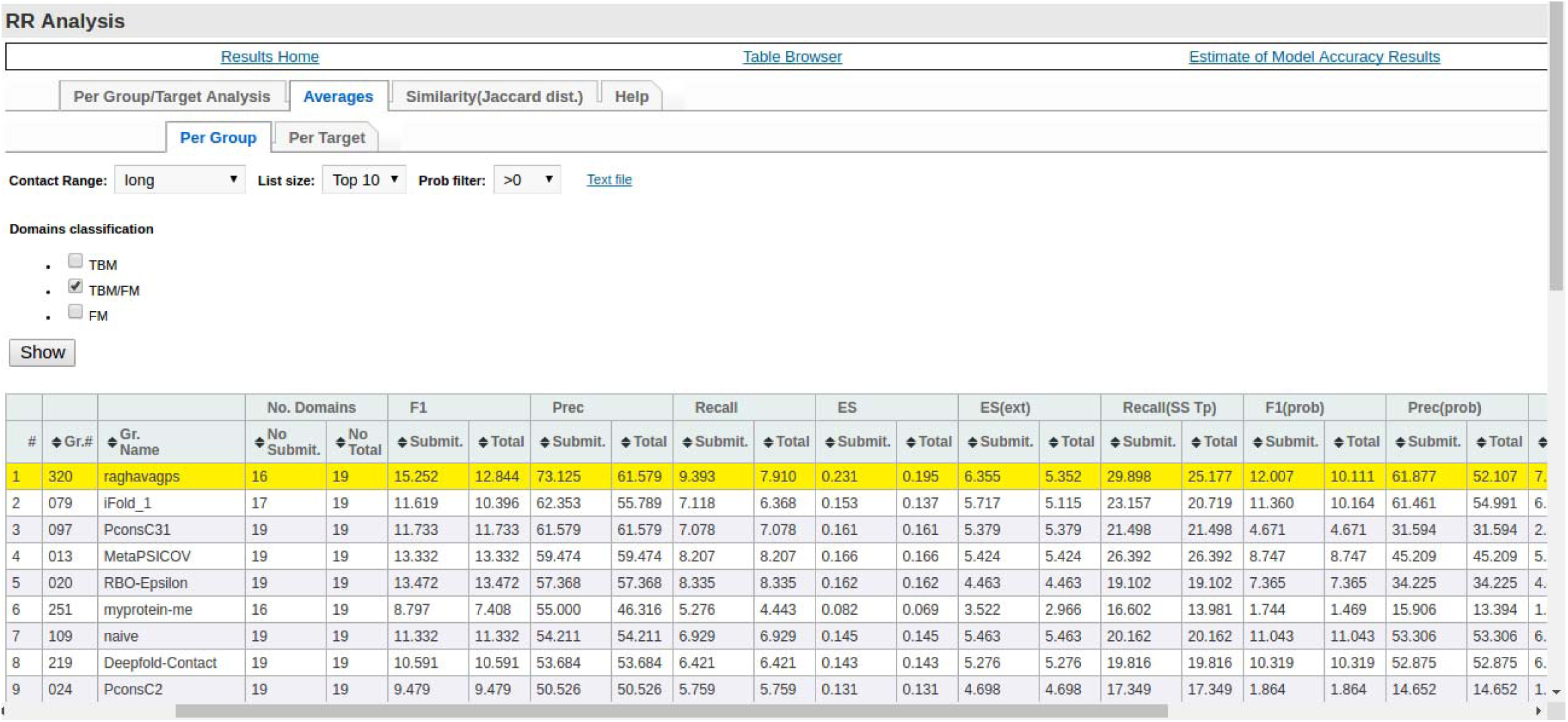
Screenshot of the performance or precision of different methods for Top10 list size and ‘long’ range contacts on CASP12 TBM/FM targets.

**Figure S5.**
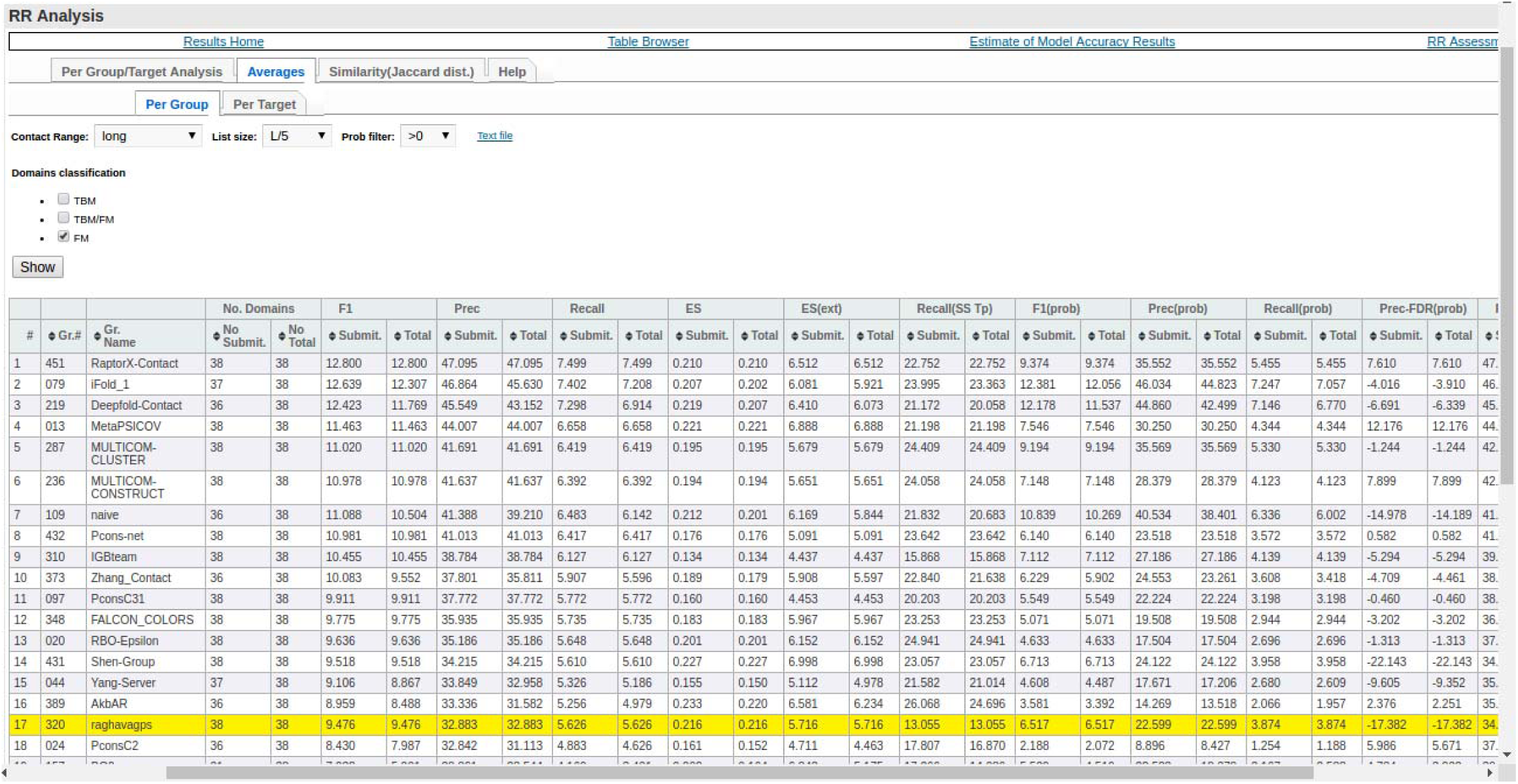
Screenshot of the performance or precision of different methods for L/5 list size and ‘long’ range contacts on CASP12 FM targets.

**Figure S6.**
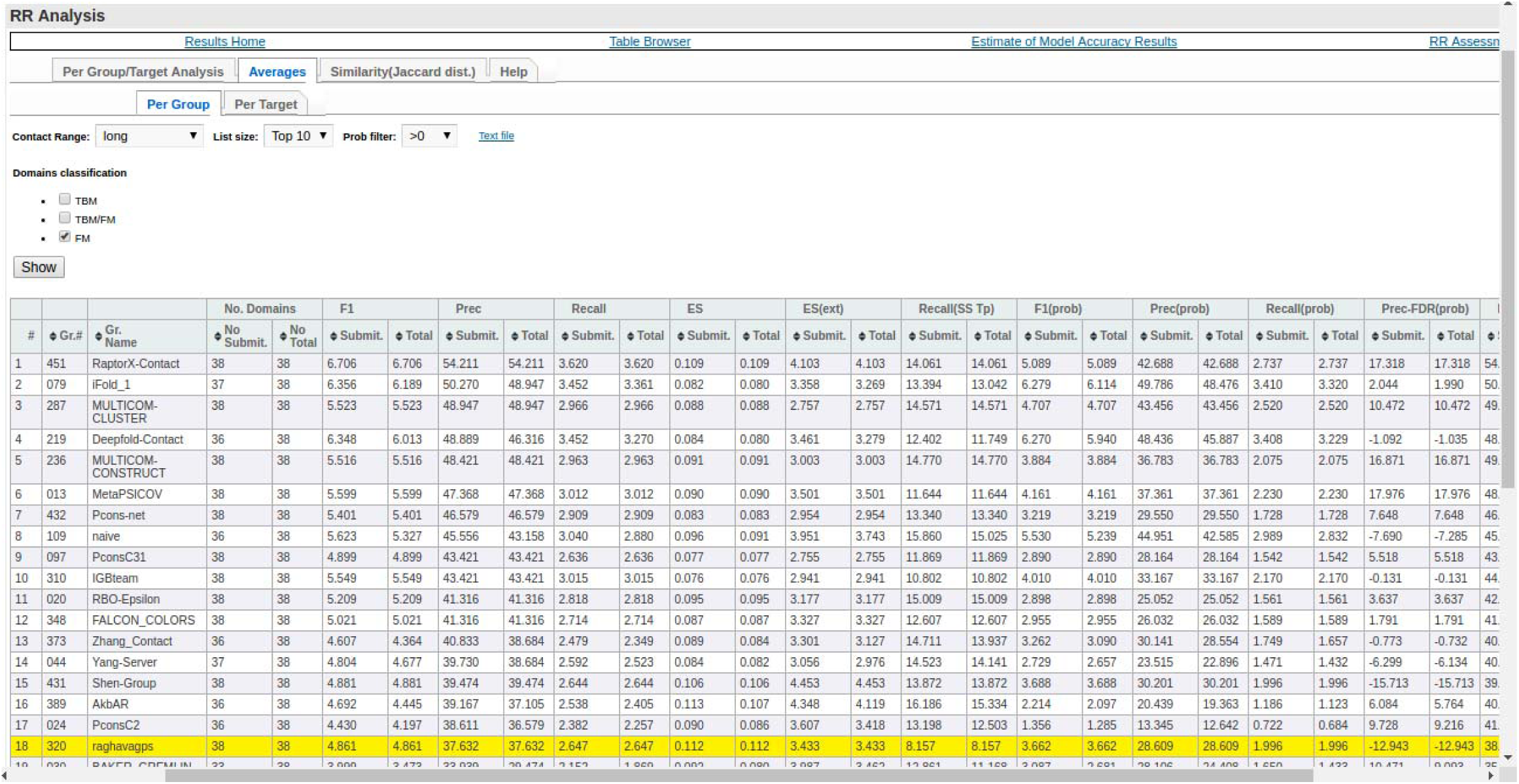
Screenshot of the performance or precision of different methods for Top10 list size and ‘long’ range contacts on CASP12 FM targets.

**Figure S7.**
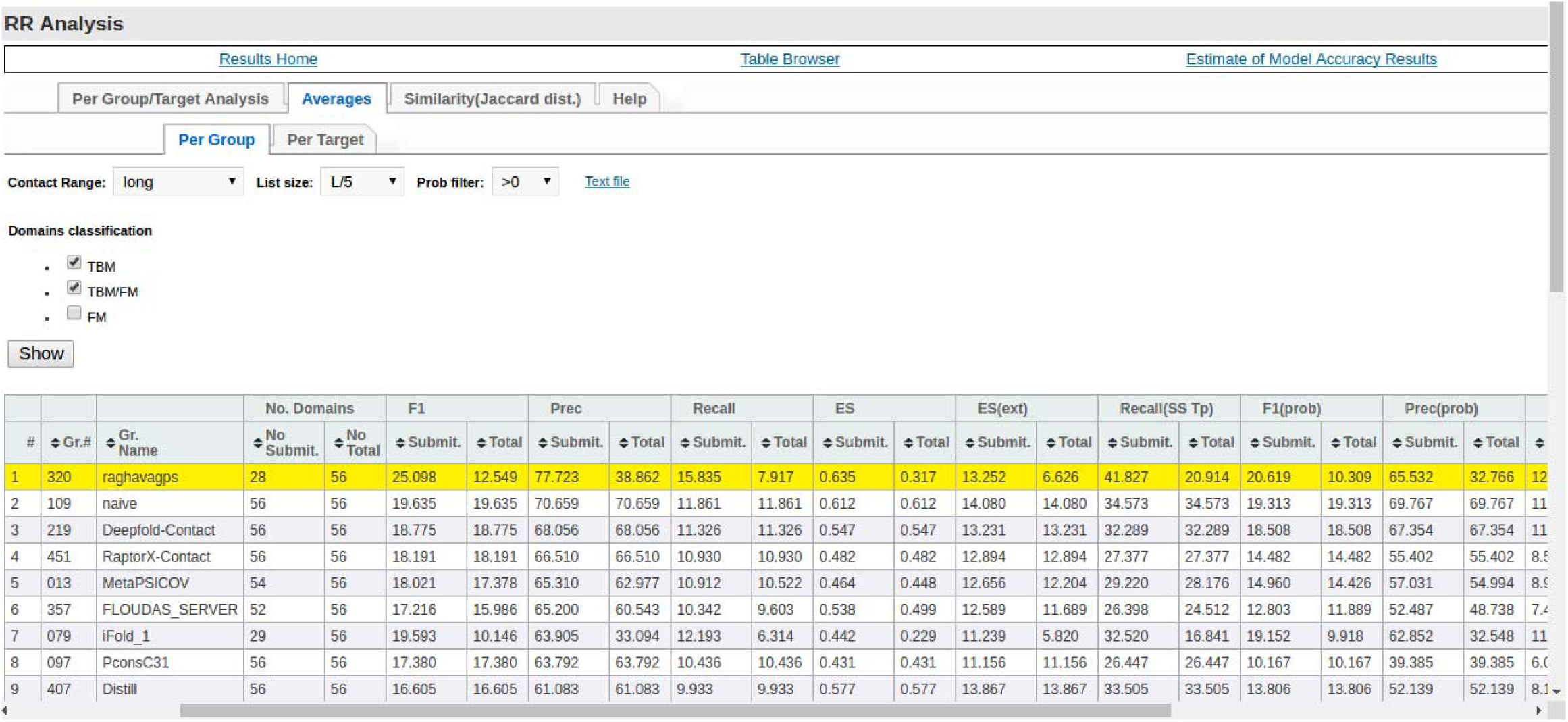
Screenshot of the performance or precision of different methods for L/5 list size and ‘long’ range contacts on CASP12 TBM and TBM/FM targets.

**Figure S8.**
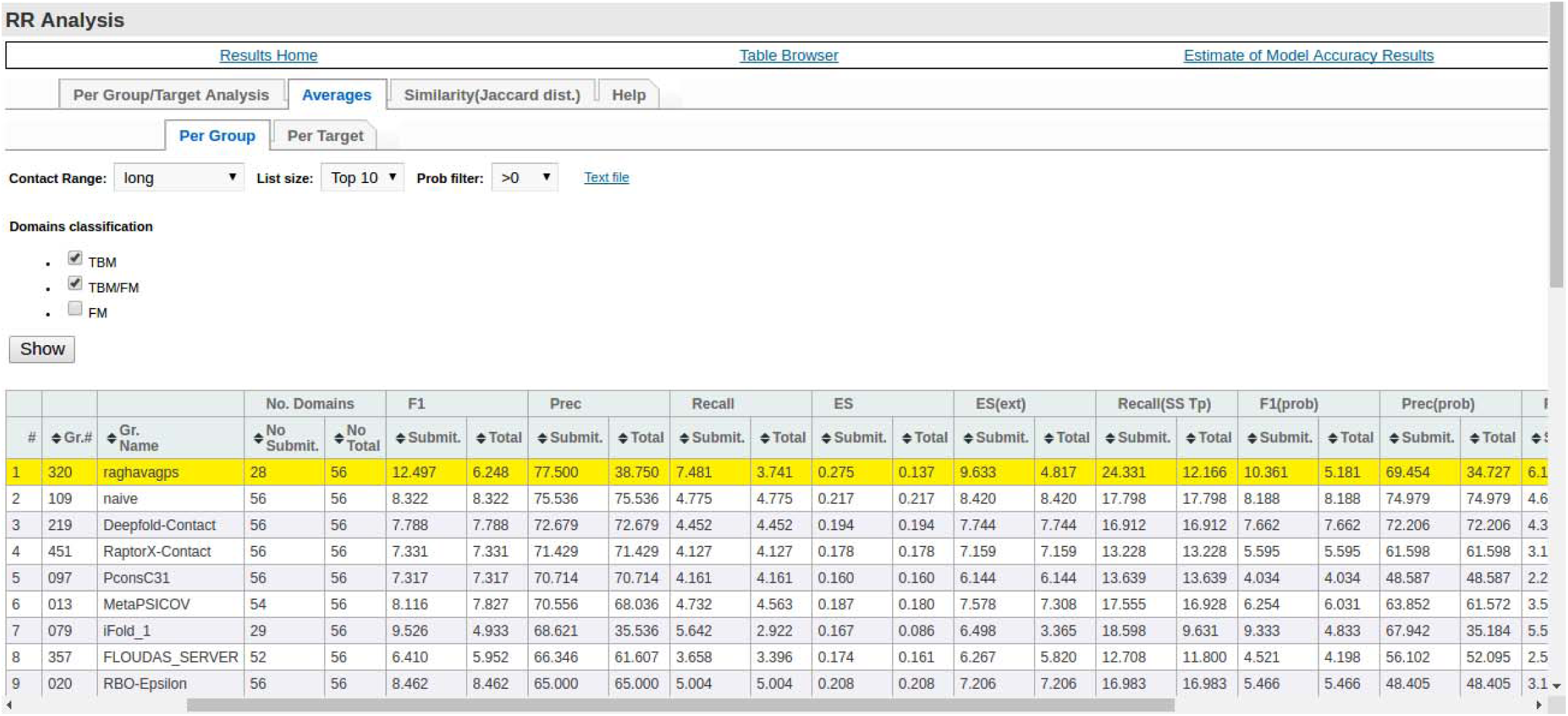
Screenshot of the performance or precision of different methods for Top10 list size and ‘long’ range contacts on CASP12 TBM and TBM/FM targets.

**Figure S9.**
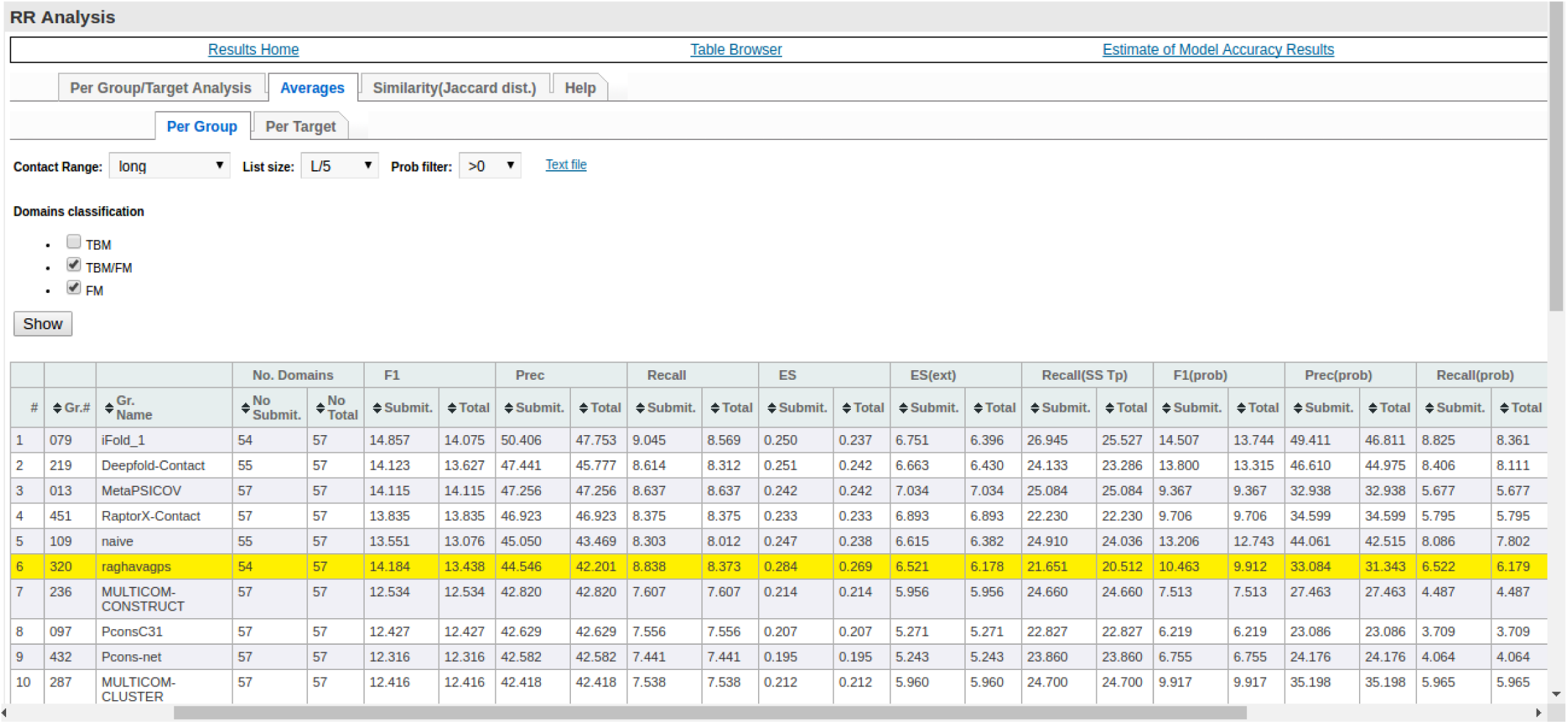
Screenshot of the performance or precision of different methods for L/5 list size and ‘long’ range contacts on CASP12 TBM/FM and FM targets.

**Figure S10.**
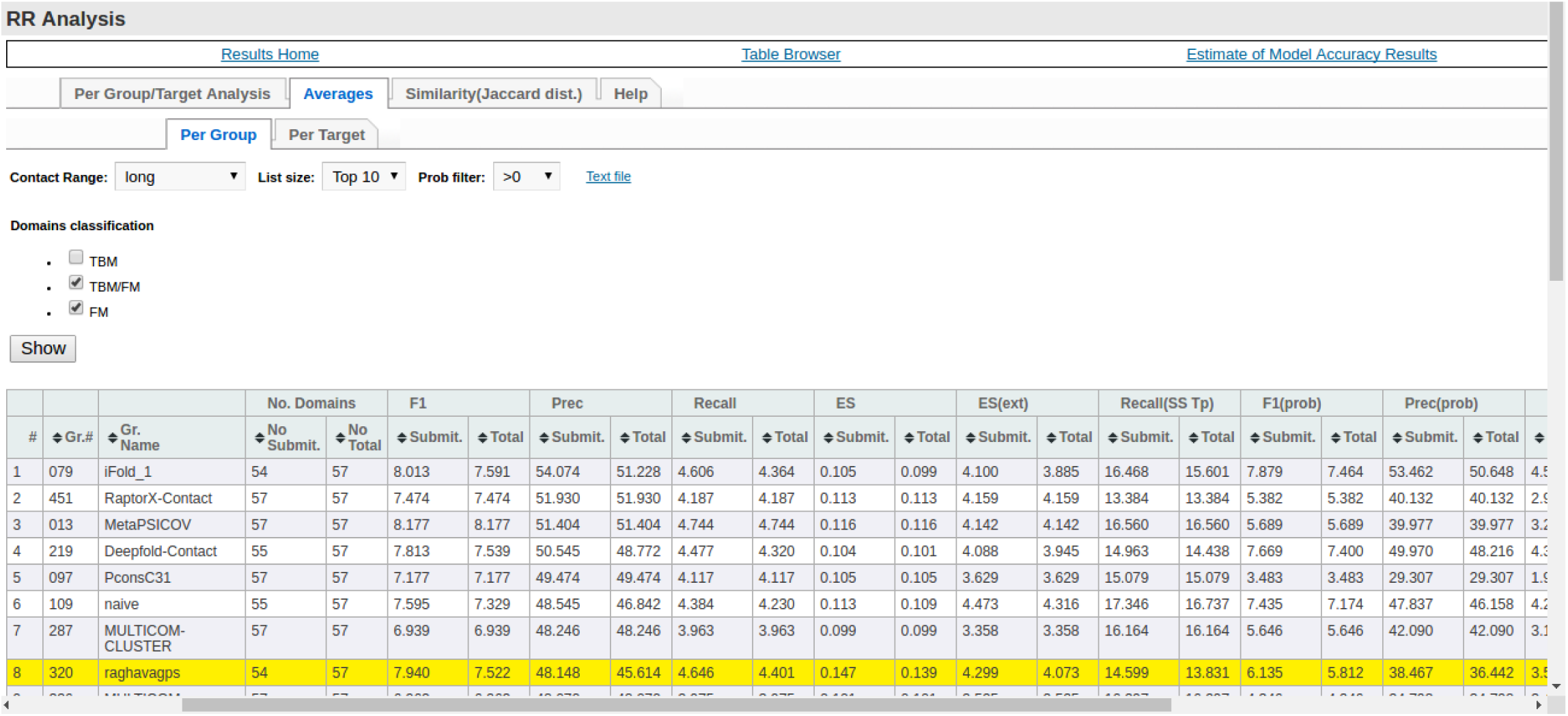
Screenshot of the performance or precision of different methods for Top10 list size and ‘long’ range contacts on CASP12 TBM/FM and FM targets.

**Figure S11.**
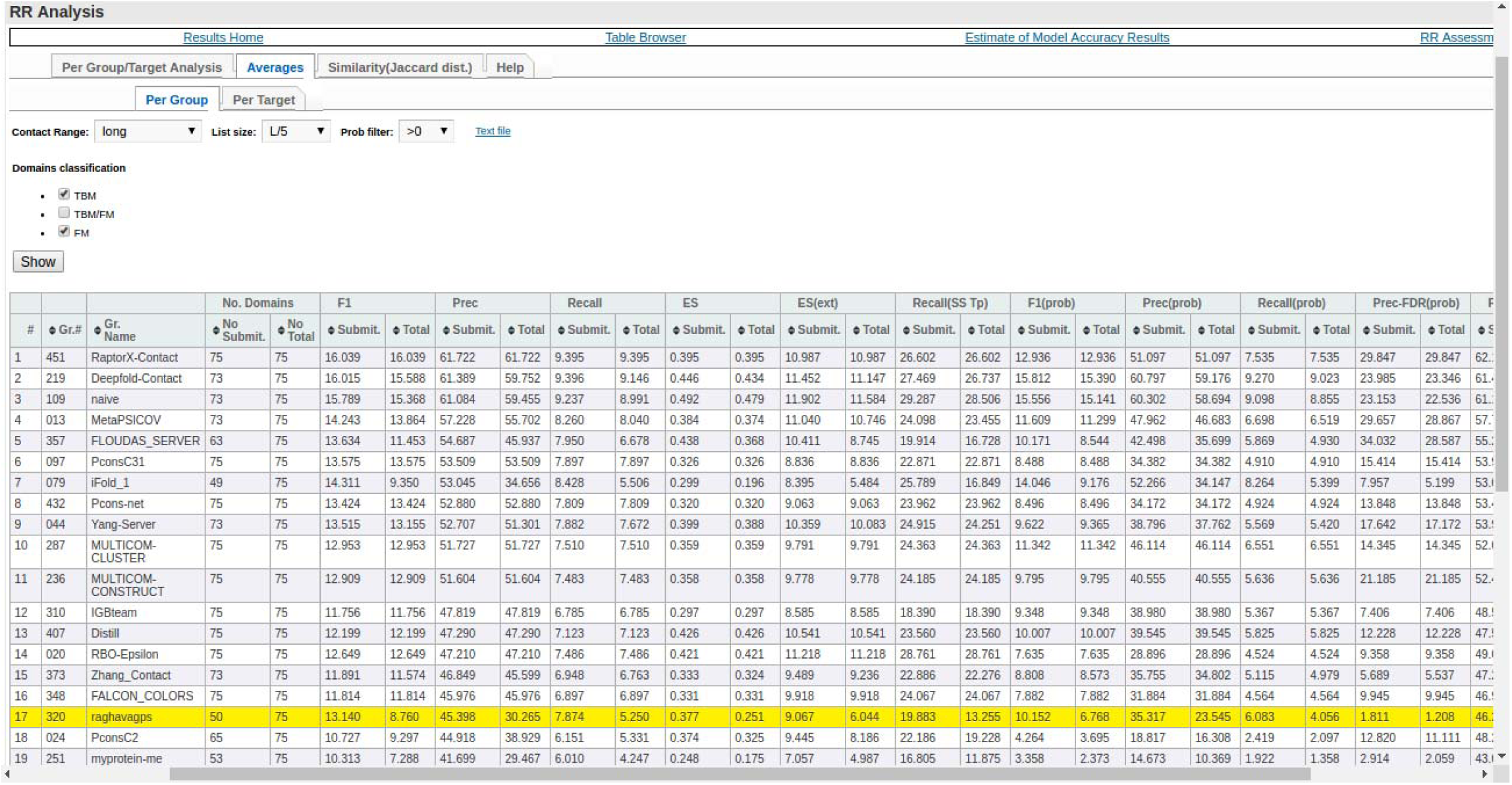
Screenshot of the performance or precision of different methods for L/5 list size and ‘long’ range contacts on CASP12 TBM and FM targets.

**Figure S12.**
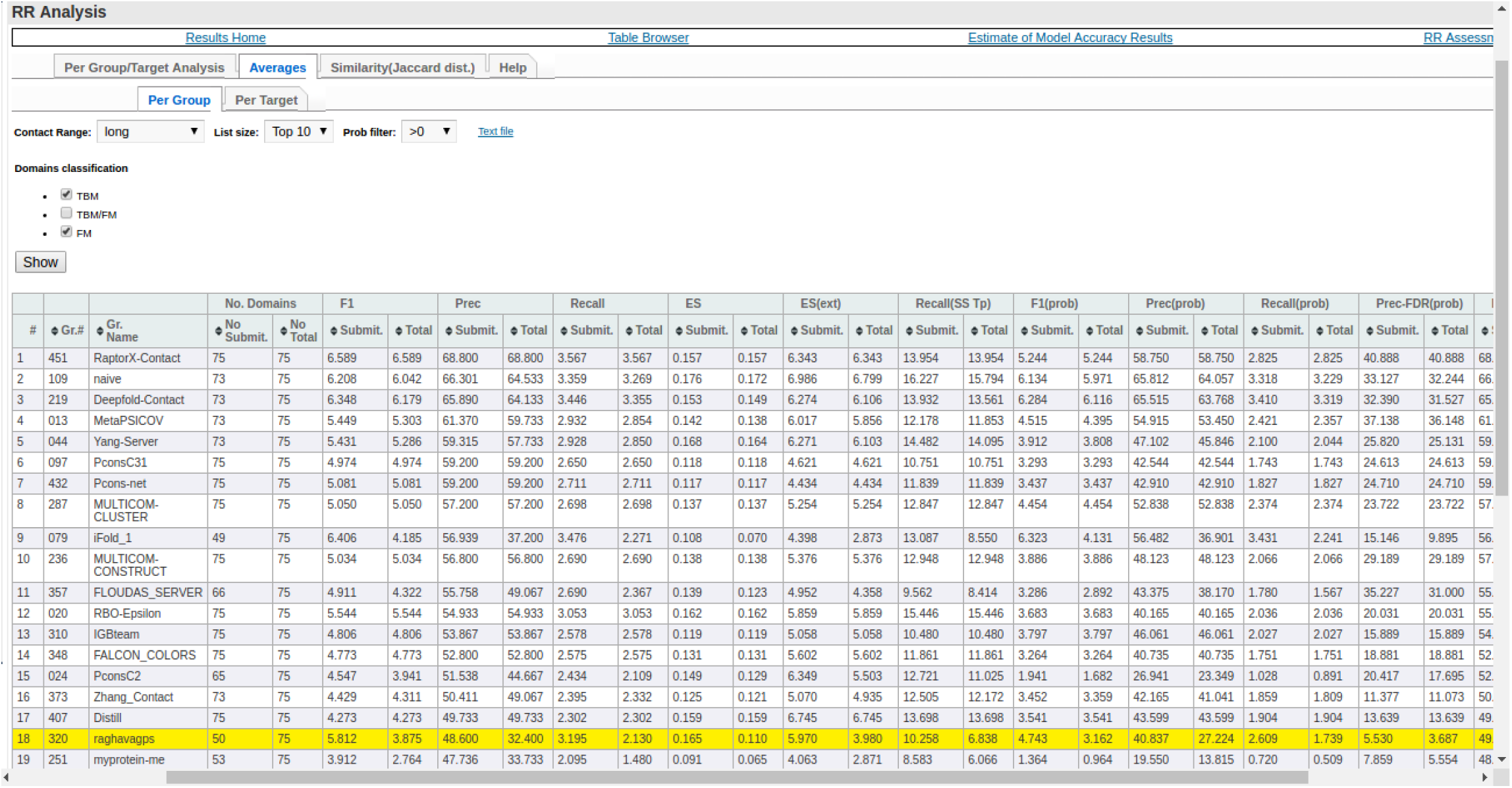
Screenshot of the performance or precision of different methods for Top10 list size and ‘long’ range contacts on CASP12 TBM and FM targets.

**Figure S13.**
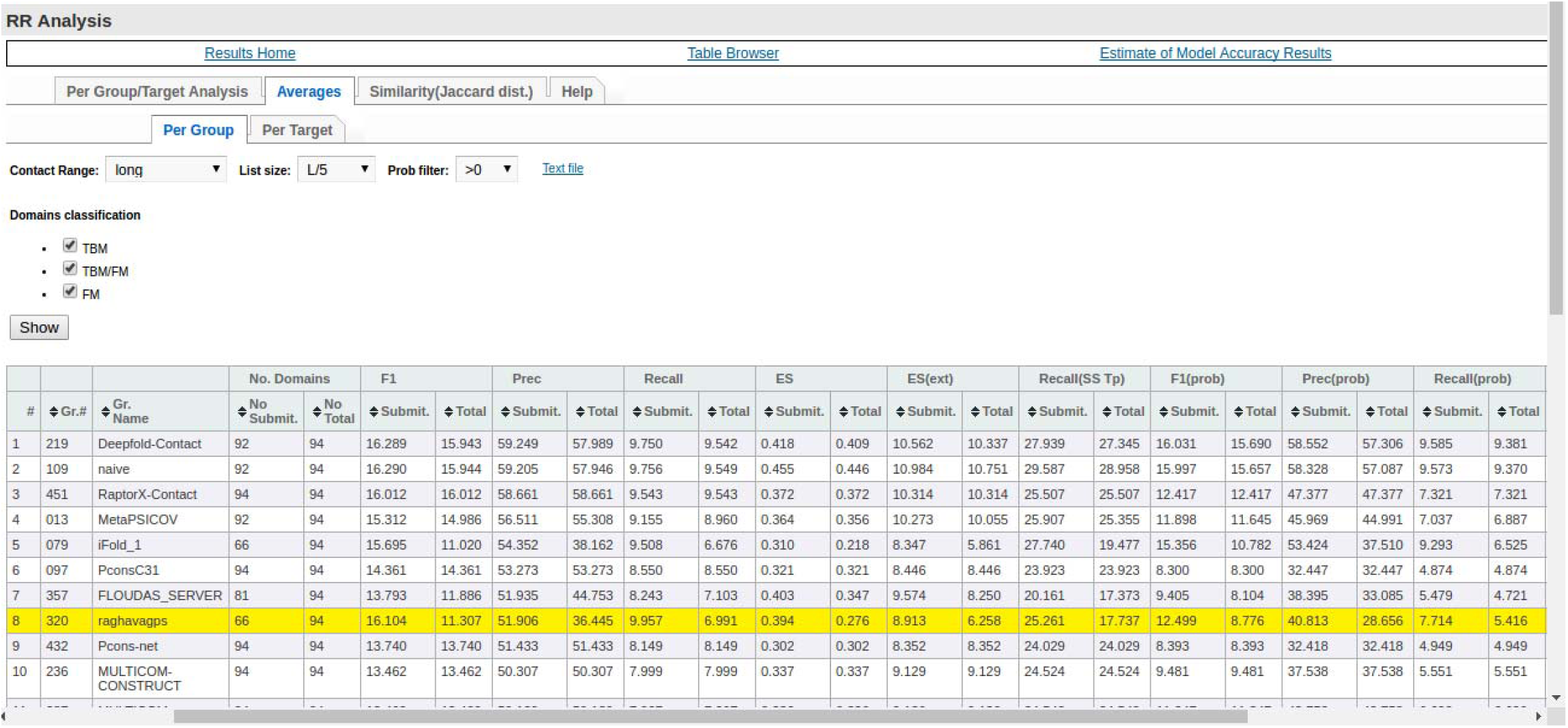
Screenshot of the performance or precision of different methods for L/5 list size and ‘long’ range contacts on CASP12 all category targets (TBM, FM and TBM/FM).

**Figure S14.**
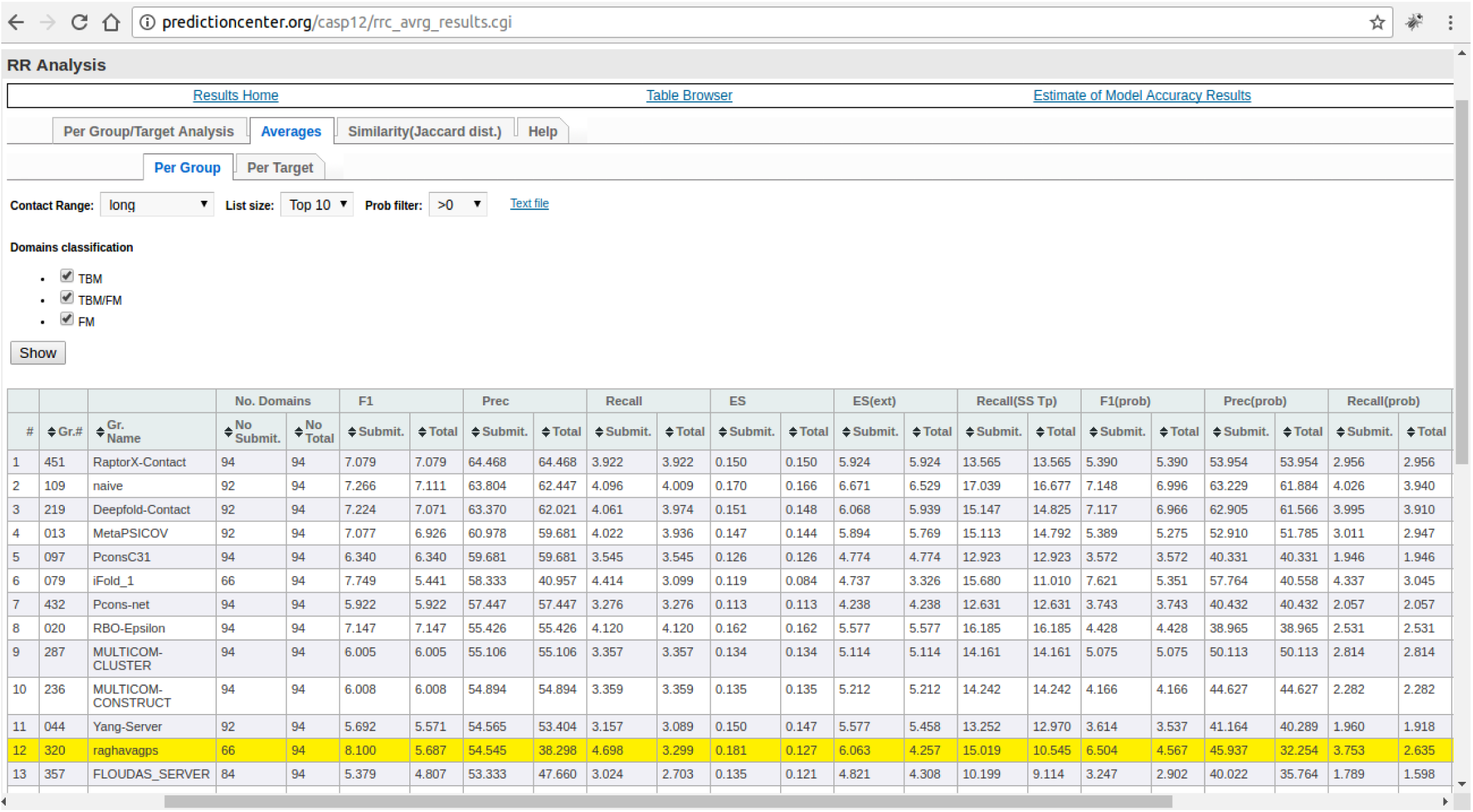
Screenshot of the performance or precision of different methods for Top10 list size and ‘long’ range contacts on CASP12 all category targets (TBM, FM and TBM/FM).

**Figure S15.**
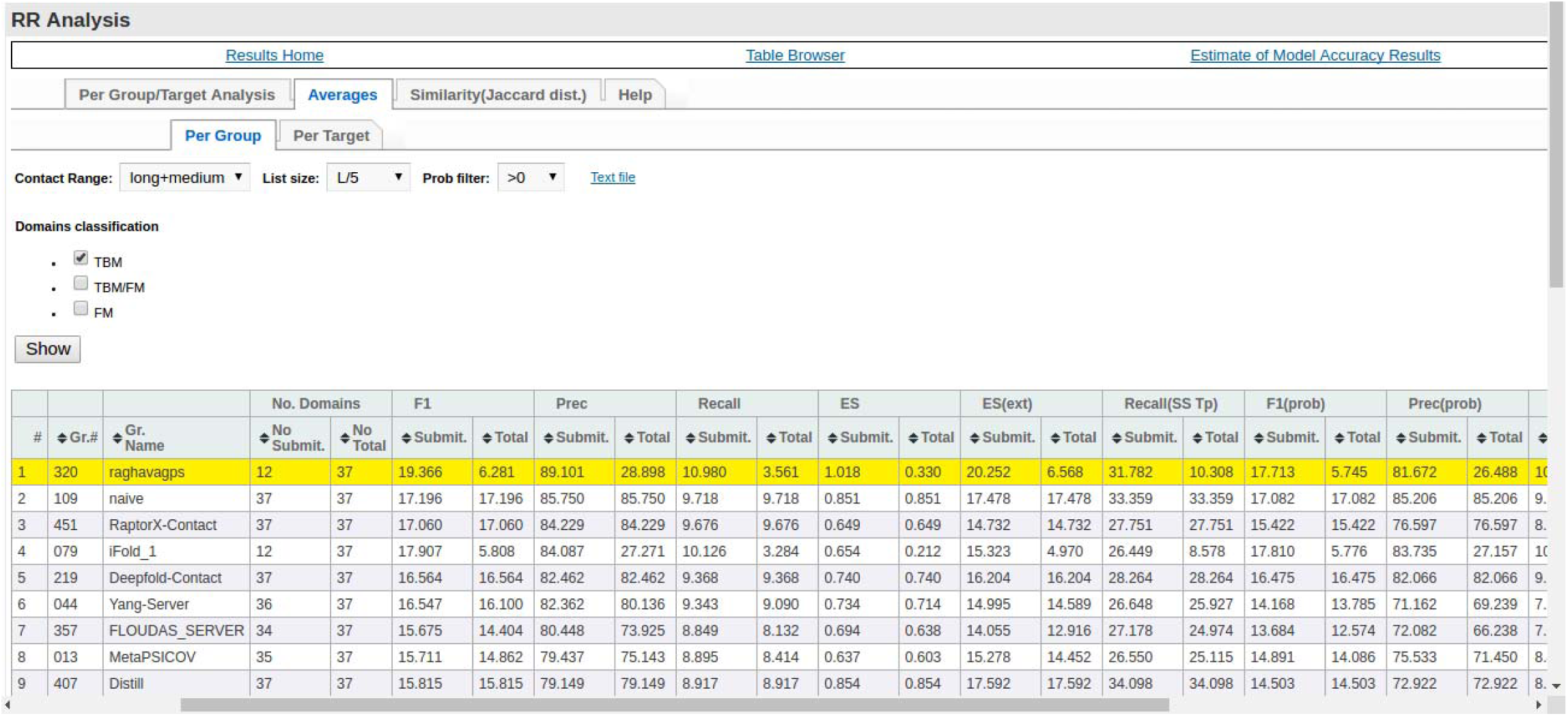
Screenshot of the performance or precision of different methods for L/5 list size and ‘long+medium’ range contacts on CASP12 TBM targets.

**Figure S16.**
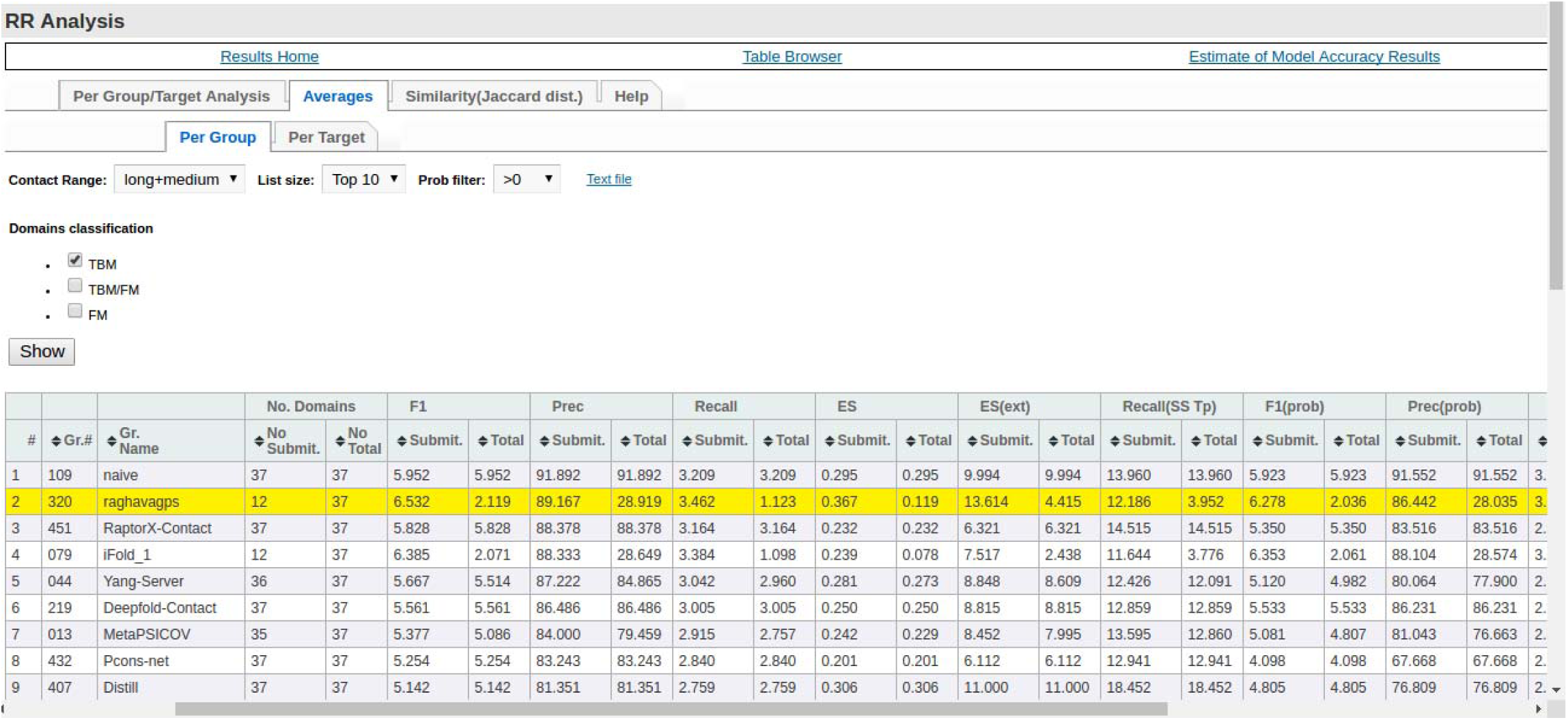
Screenshot of the performance or precision of different methods for Top10 list size and ‘long+medium’ range contacts on CASP12 TBM targets.

**Figure S17.**
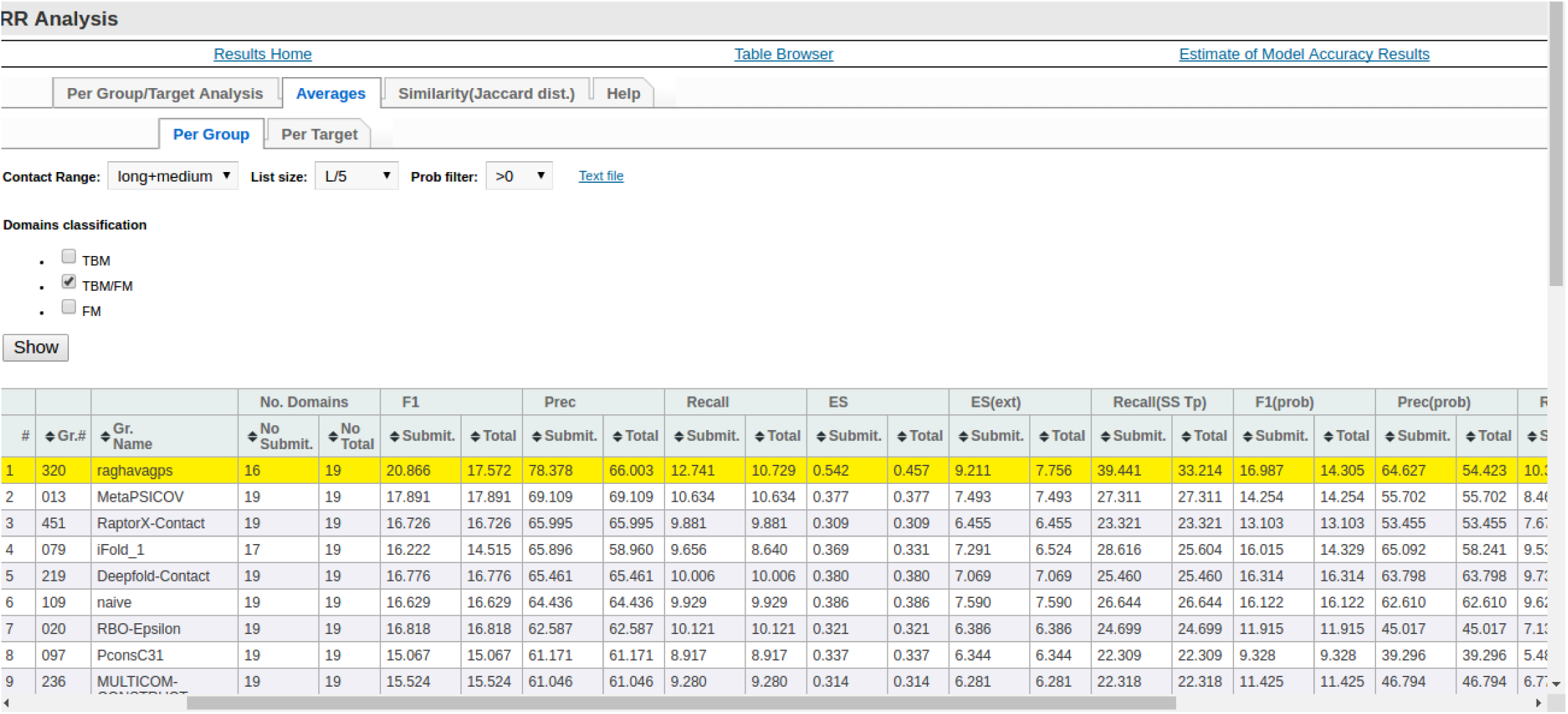
Screenshot of the performance or precision of different methods for L/5 list size and ‘long+medium’ range contacts on CASP12 TBM/FM targets.

**Figure S18.**
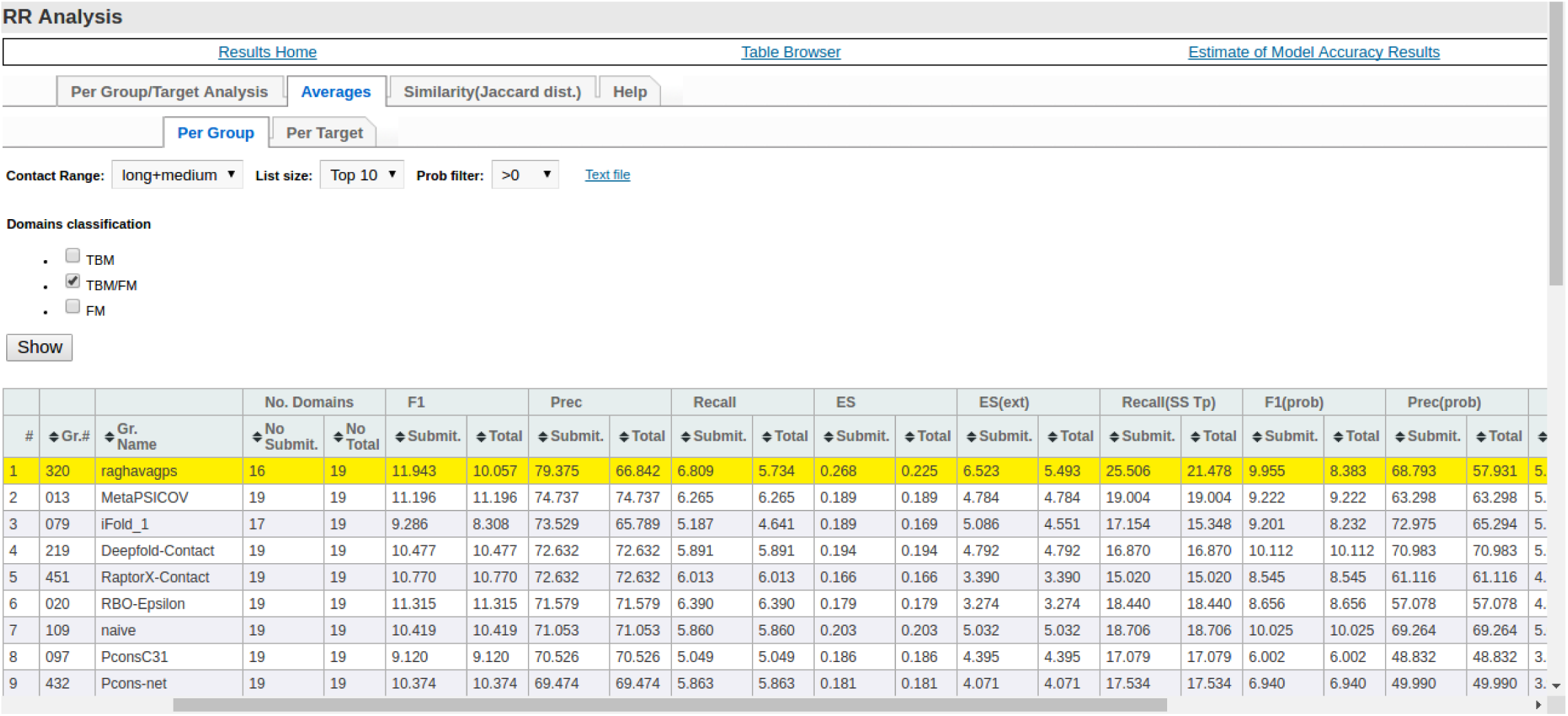
Screenshot of the performance or precision of different methods for Top10 list size and ‘long+medium’ range contacts on CASP12 TBM/FM targets.

**Figure S19.**
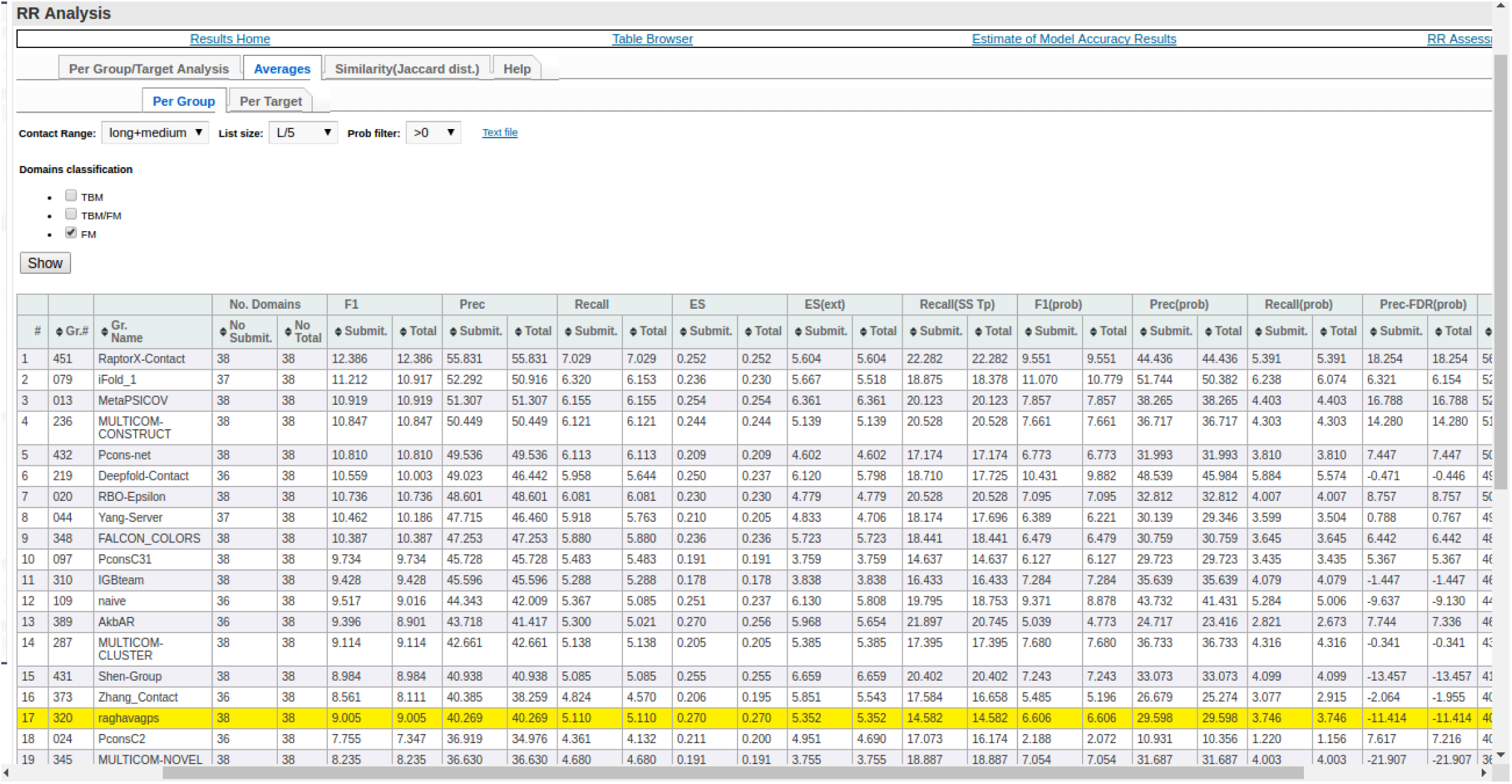
Screenshot of the performance or precision of different methods for L/5 list size and ‘long+medium’ range contacts on CASP12 FM targets.

**Figure S20.**
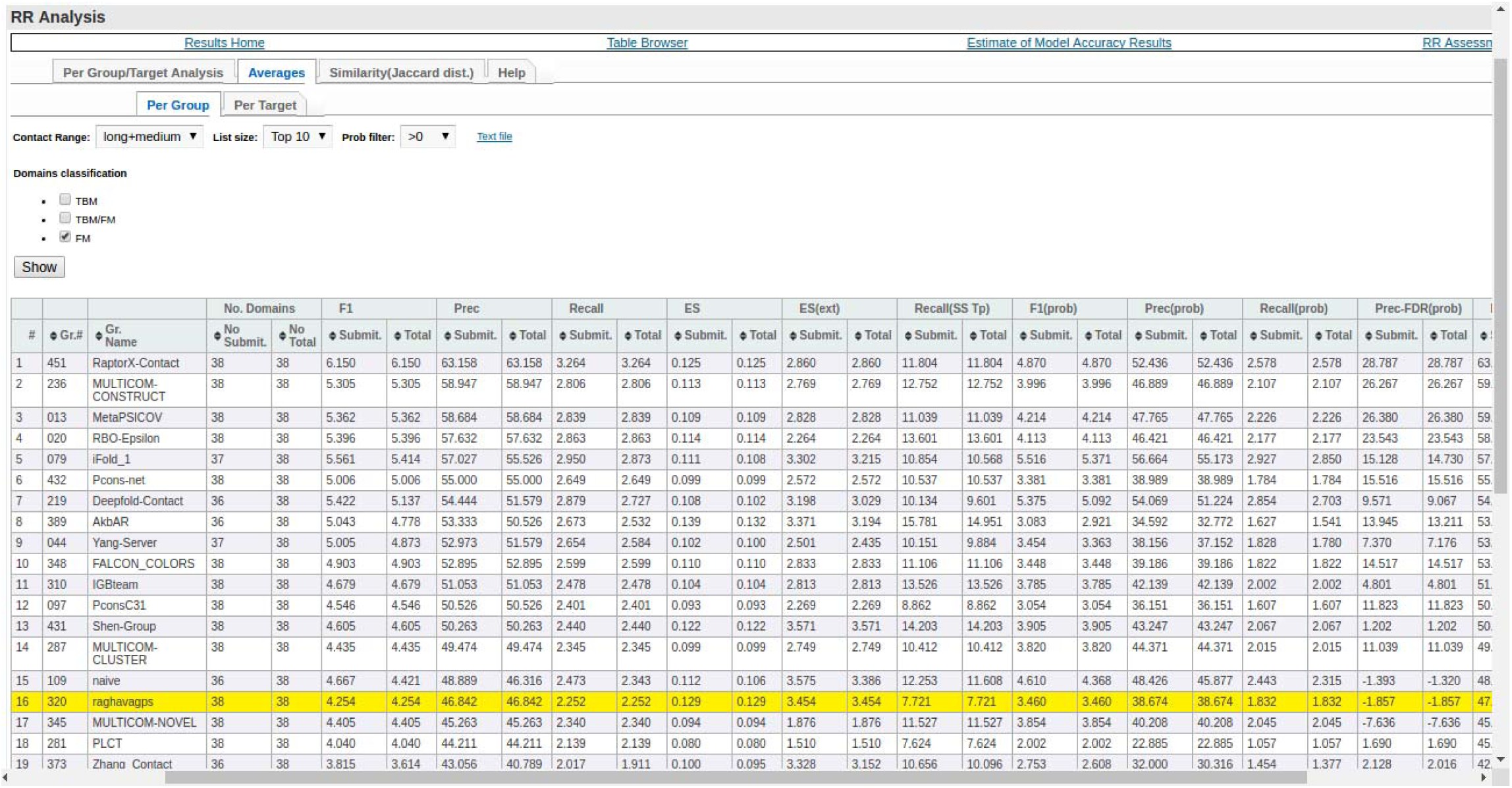
Screenshot of the performance or precision of different methods for Top10 list size and ‘long+medium’ range contacts on CASP12 FM targets.

**Figure S21.**
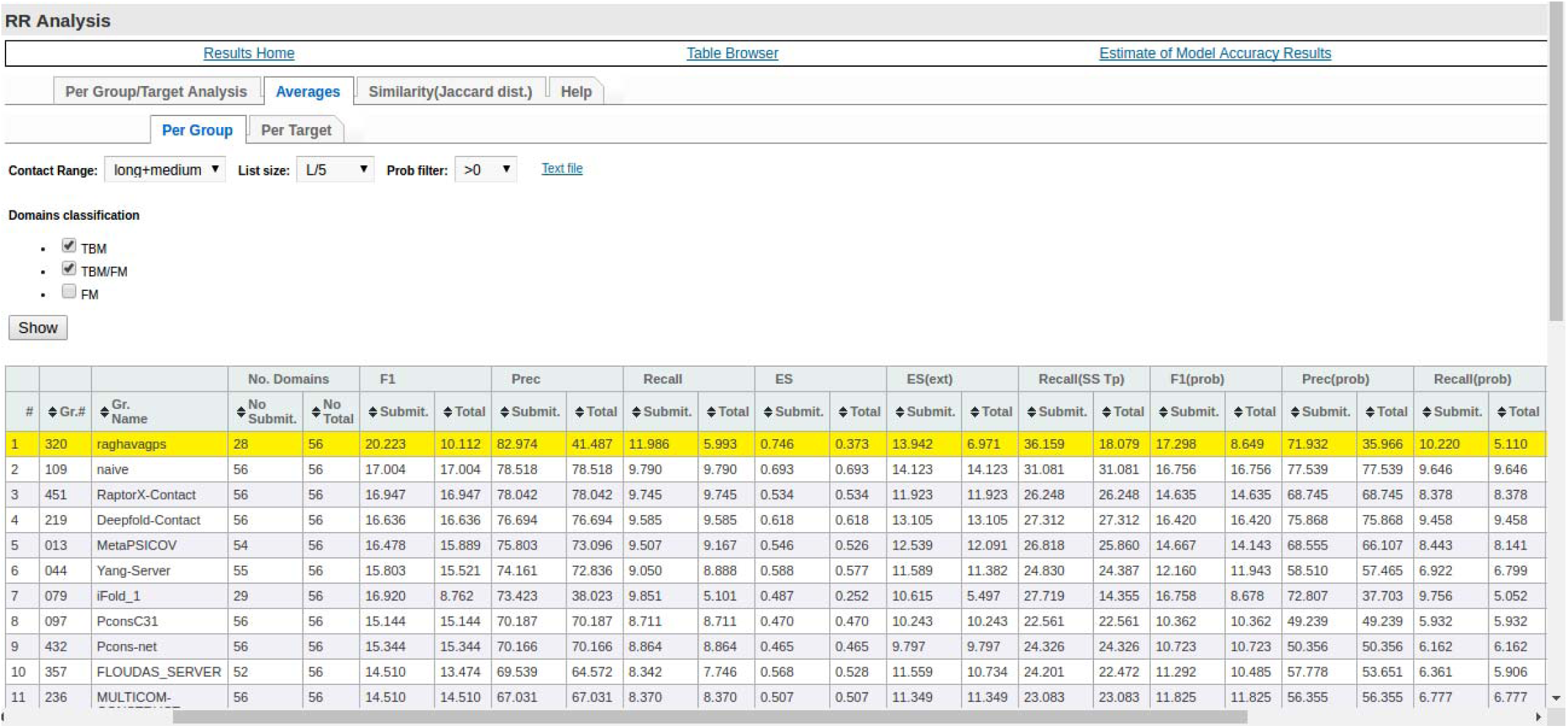
Screenshot of the performance or precision of different methods for L/5 list size and ‘long+medium’ range contacts on CASP12 TBM and TBM/FM targets.

**Figure S22.**
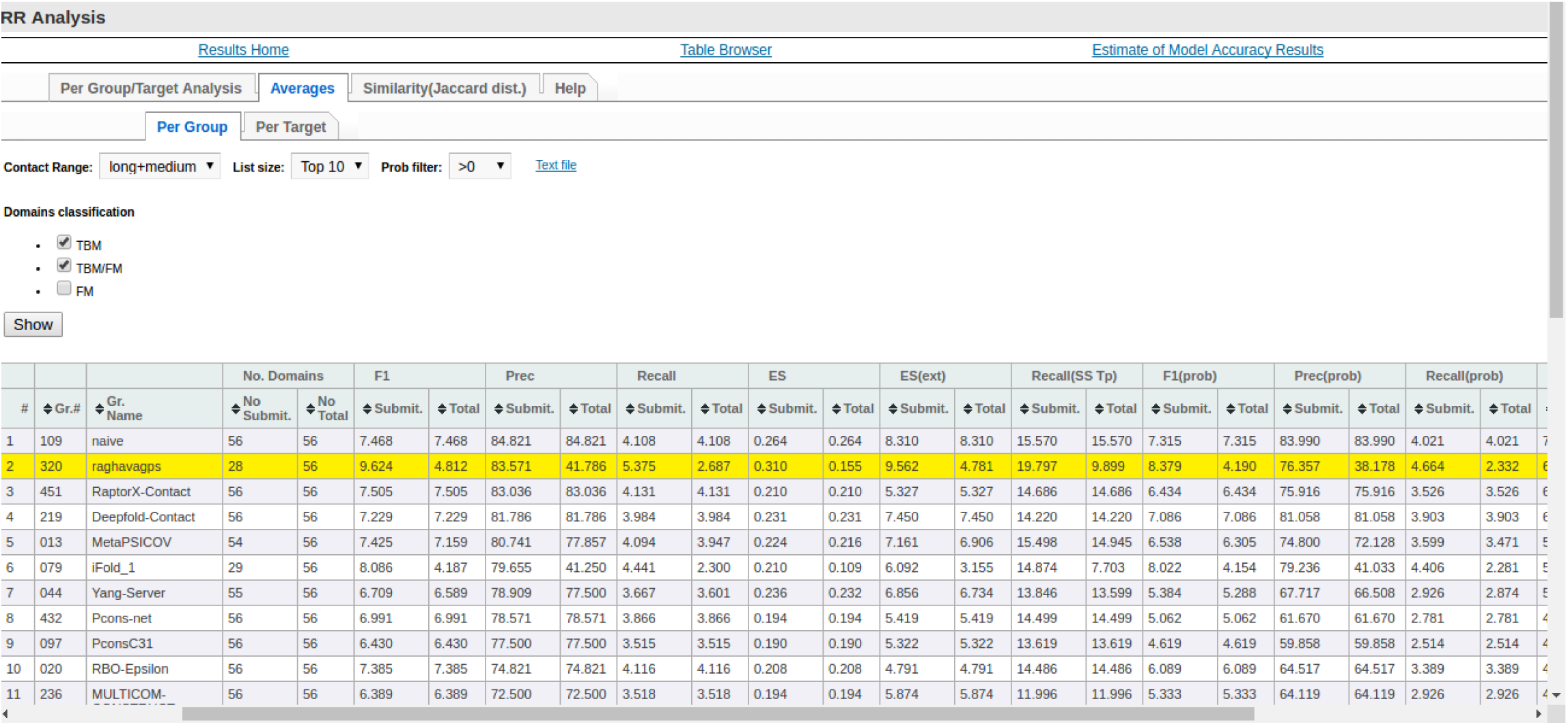
Screenshot of the performance or precision of different methods for Top10 list size and ‘long+medium’ range contacts on CASP12 TBM and TBM/FM targets.

**Figure S23.**
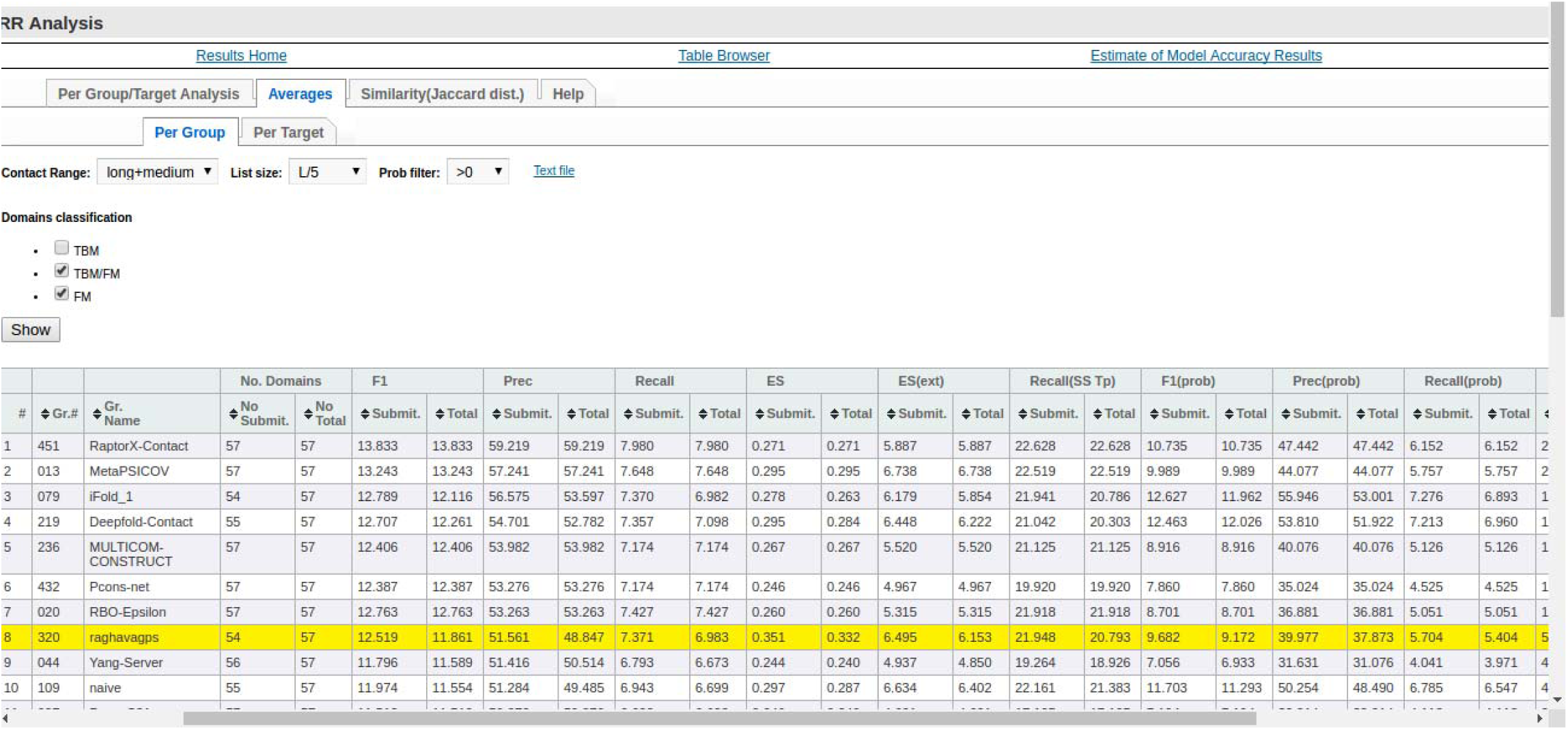
Screenshot of the performance or precision of different methods for L/5 list size and ‘long+medium’ range contacts on CASP12 TBM/FM and FM targets.

**Figure S24.**
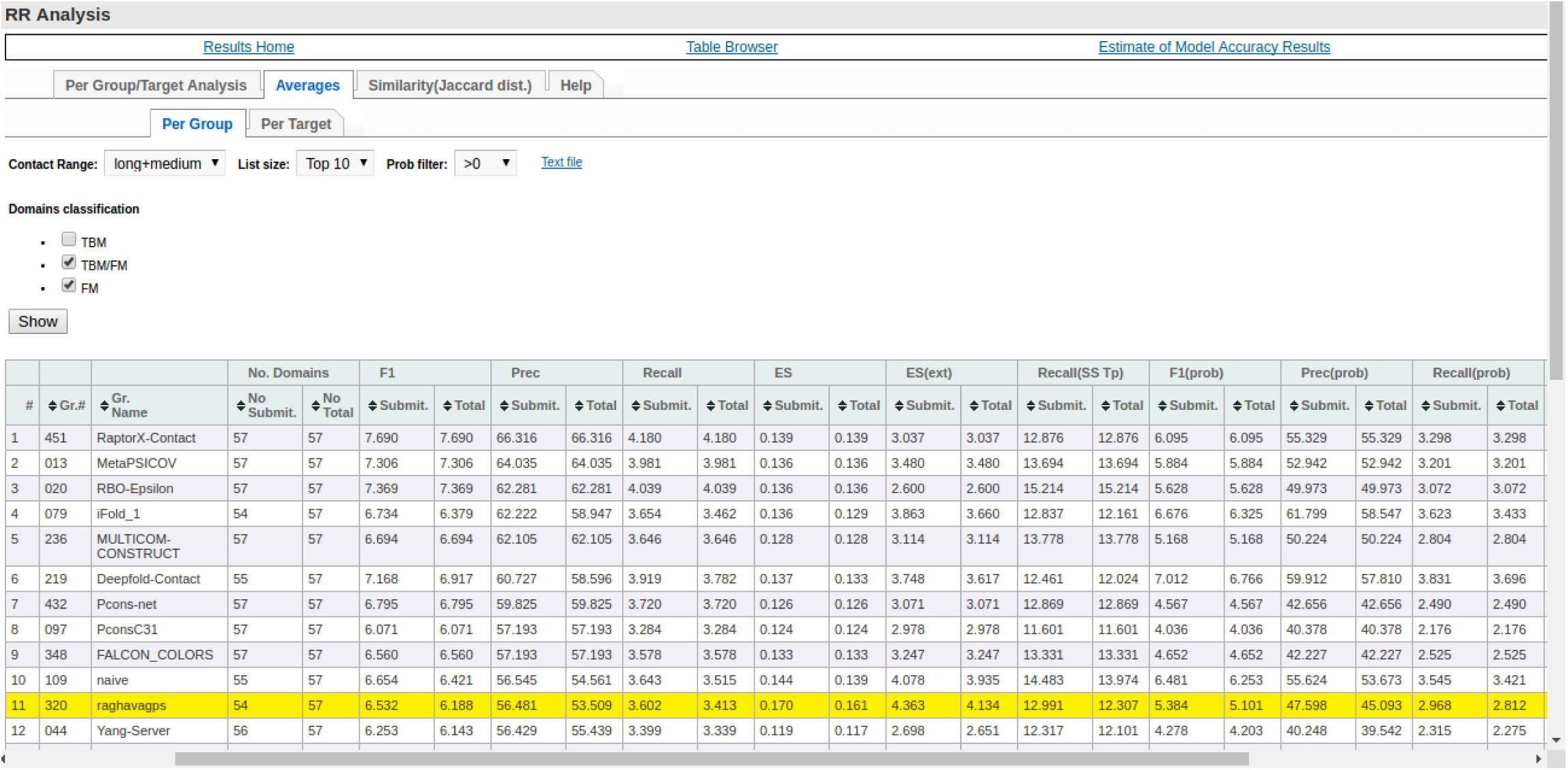
Screenshot of the performance or precision of different methods for Top10 list size and ‘long+medium’ range contacts on CASP12 TBM/FM and FM targets.

**Figure S25.**
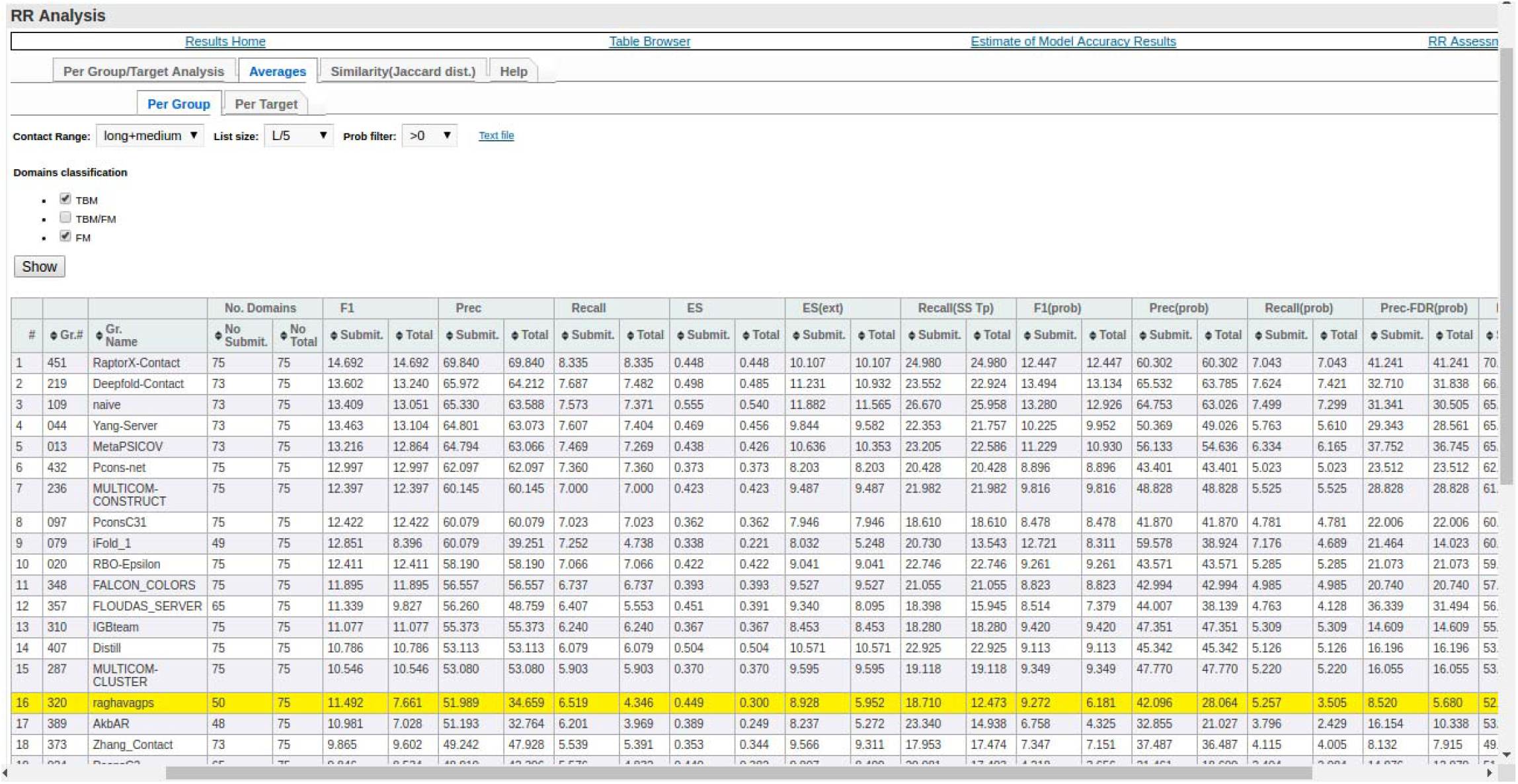
Screenshot of the performance or precision of different methods for L/5 list size and ‘long+medium’ range contacts on CASP12 TBM and FM targets.

**Figure S26.**
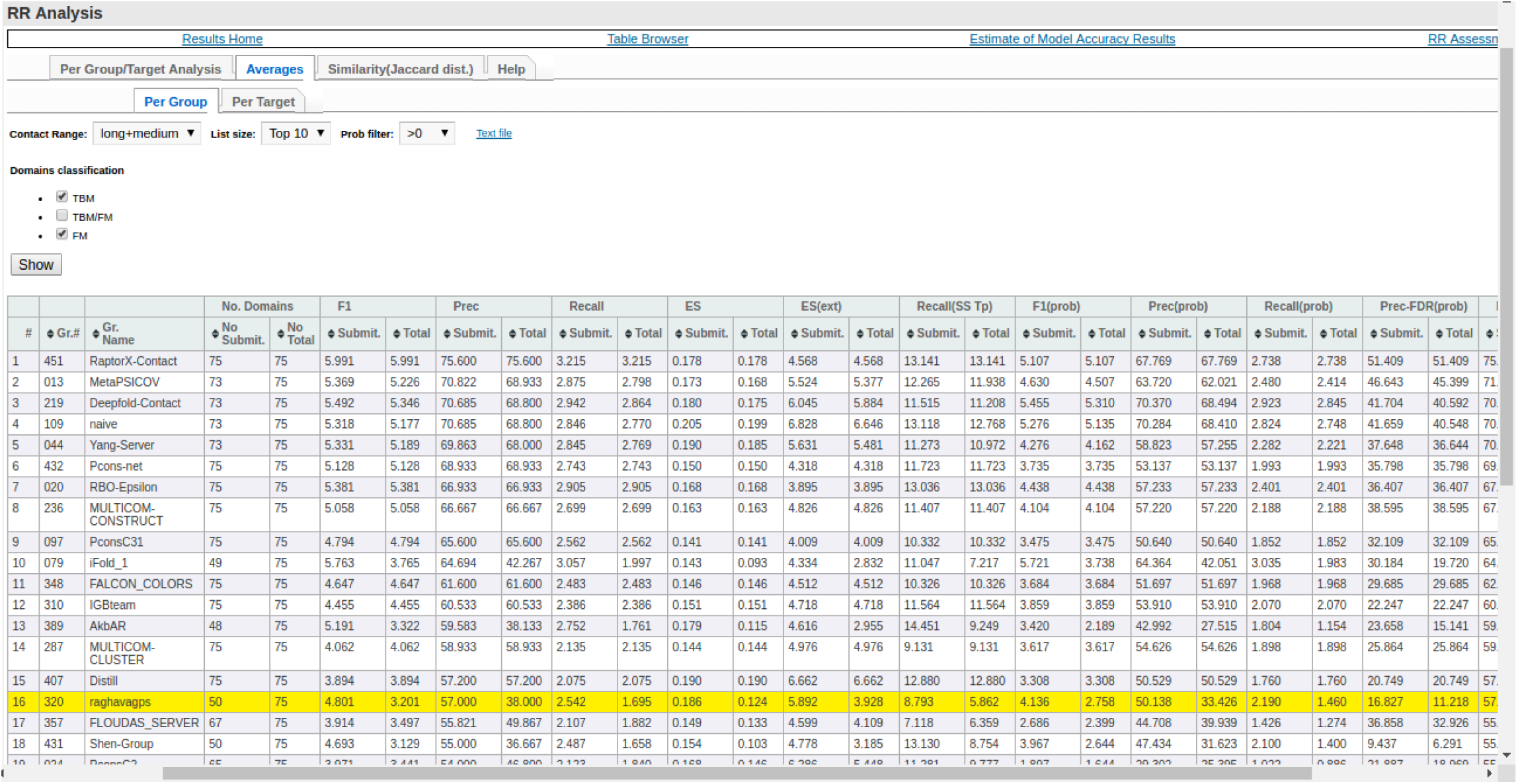
Screenshot of the performance or precision of different methods for Top10 list size and ‘long+medium’ range contacts on CASP12 TBM and FM targets.

**Figure S27.**
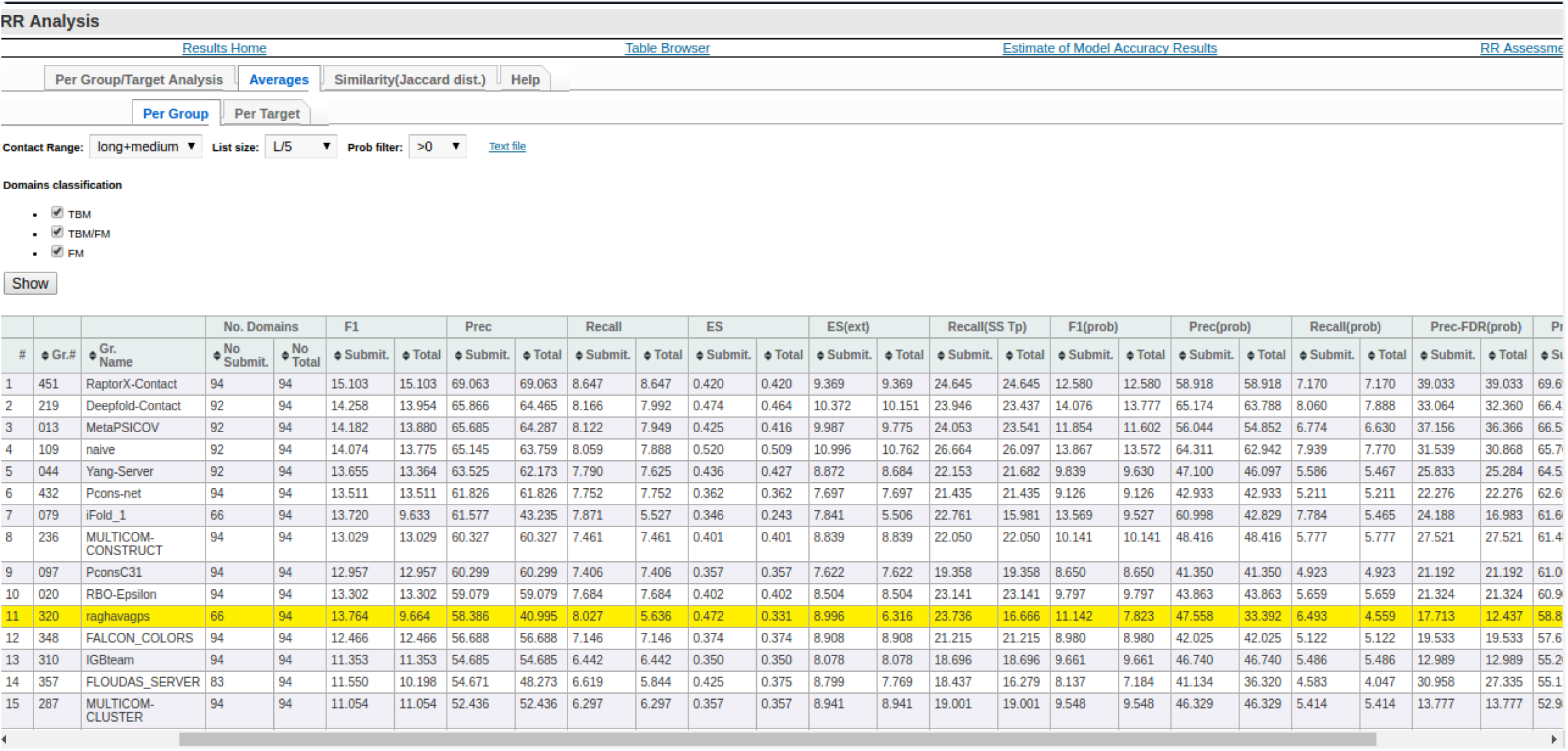
Screenshot of the performance or precision of different methods for L/5 list size and ‘long+medium’ range contacts on CASP12 all category targets (TBM, FM and TBM/FM).

**Figure S28.**
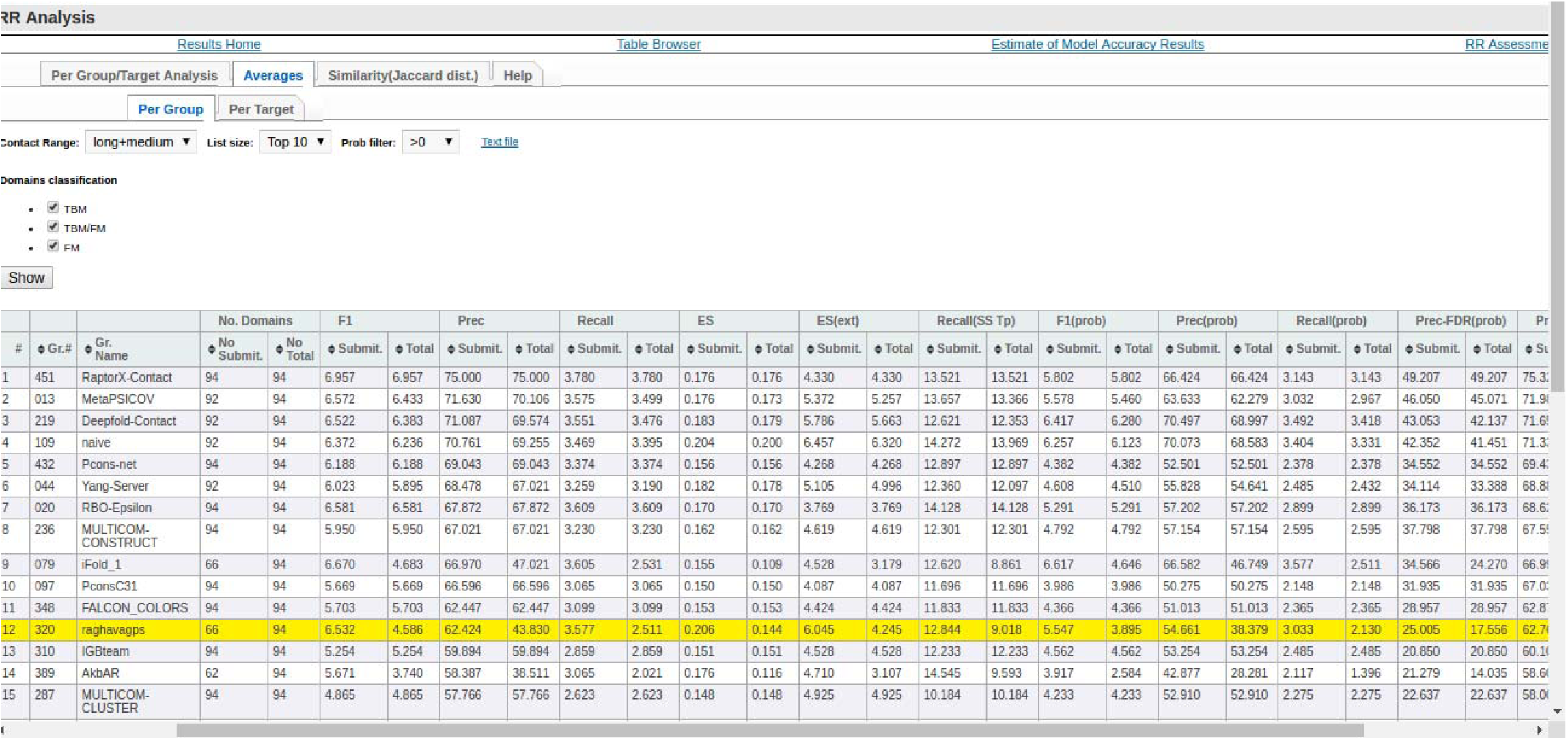
Screenshot of the performance or precision of different methods for Top10 list size and ‘long+medium’ range contacts on CASP12 all category targets (TBM, FM and TBM/FM).

**Supplementary Table S1:**
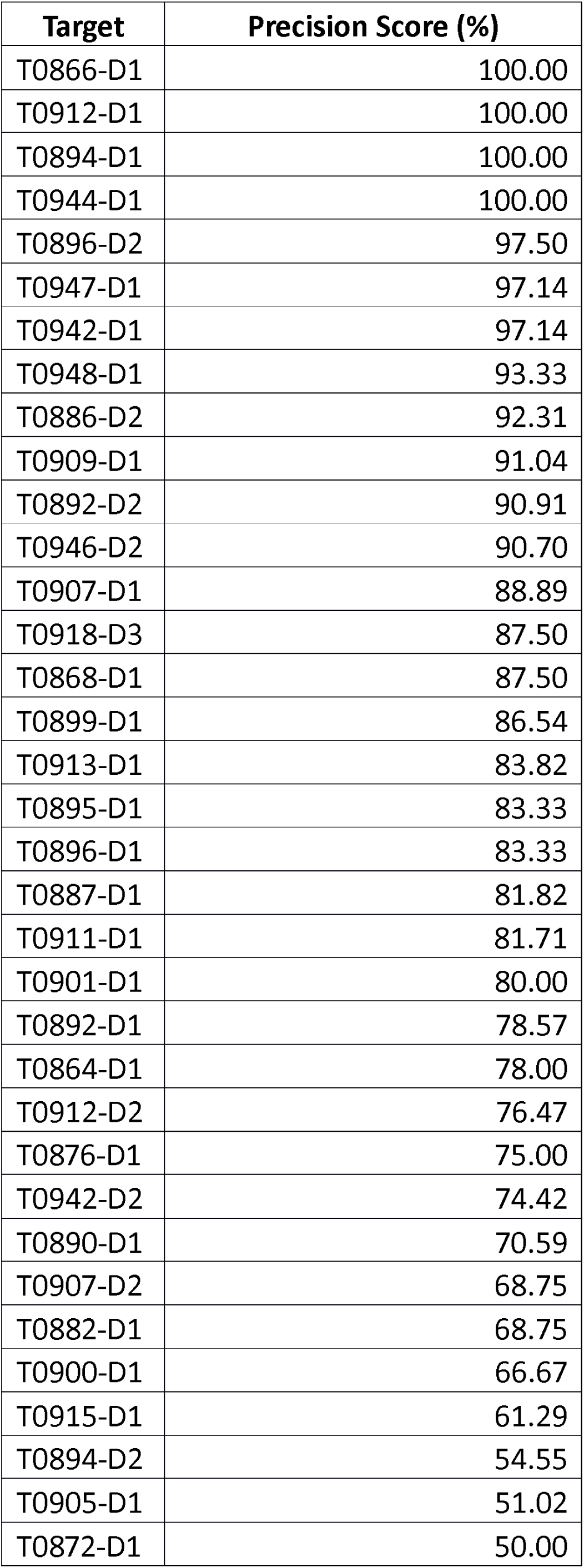
List of targets with average precision value 50% or above when list size is L/5 and contact range is Long.

| **Target** | **Precision Score (%)** |
| --- | --- |
| T0866-D1 | 100.00 |
| T0912-D1 | 100.00 |
| T0894-D1 | 100.00 |
| T0944-D1 | 100.00 |
| T0896-D2 | 97.50 |
| T0947-D1 | 97.14 |
| T0942-D1 | 97.14 |
| T0948-D1 | 93.33 |
| T0886-D2 | 92.31 |
| T0909-D1 | 91.04 |
| T0892-D2 | 90.91 |
| T0946-D2 | 90.70 |
| T0907-D1 | 88.89 |
| T0918-D3 | 87.50 |
| T0868-D1 | 87.50 |
| T0899-D1 | 86.54 |
| T0913-D1 | 83.82 |
| T0895-D1 | 83.33 |
| T0896-D1 | 83.33 |
| T0887-D1 | 81.82 |
| T0911-D1 | 81.71 |
| T0901-D1 | 80.00 |
| T0892-D1 | 78.57 |
| T0864-D1 | 78.00 |
| T0912-D2 | 76.47 |
| T0876-D1 | 75.00 |
| T0942-D2 | 74.42 |
| T0890-D1 | 70.59 |
| T0907-D2 | 68.75 |
| T0882-D1 | 68.75 |
| T0900-D1 | 66.67 |
| T0915-D1 | 61.29 |
| T0894-D2 | 54.55 |
| T0905-D1 | 51.02 |
| T0872-D1 | 50.00 |

**Supplementary Table S2:** List of targets with average precision value 50% or above when list size is Top10 and contact range is Long.

| Target | Precision Score (%) |
| --- | --- |
| T0896-D1 | 100.00 |
| T0866-D1 | 100.00 |
| T0946-D2 | 100.00 |
| T0944-D1 | 100.00 |
| T0942-D1 | 100.00 |
| T0918-D3 | 100.00 |
| T0886-D2 | 100.00 |
| T0912-D1 | 100.00 |
| T0892-D2 | 100.00 |
| T0894-D1 | 100.00 |
| T0901-D1 | 90.00 |
| T0912-D2 | 90.00 |
| T0896-D2 | 90.00 |
| T0907-D1 | 90.00 |
| T0899-D1 | 90.00 |
| T0868-D1 | 90.00 |
| T0948-D1 | 90.00 |
| T0909-D1 | 90.00 |
| T0892-D1 | 90.00 |
| T0947-D1 | 90.00 |
| T0913-D1 | 90.00 |
| T0911-D1 | 80.00 |
| T0915-D1 | 80.00 |
| T0864-D1 | 80.00 |
| T0882-D1 | 80.00 |
| T0907-D2 | 80.00 |
| T0895-D1 | 80.00 |
| T0904-D1 | 70.00 |
| T0918-D1 | 70.00 |
| T0894-D2 | 60.00 |
| T0890-D1 | 60.00 |
| T0888-D1 | 60.00 |
| T0898-D1 | 60.00 |
| T0887-D1 | 60.00 |
| T0876-D1 | 60.00 |
| T0900-D1 | 60.00 |
| T0898-D2 | 50.00 |
| T0870-D1 | 50.00 |
| T0942-D2 | 50.00 |

**Table S3.** List of targets with average precision value 50% or above when list size is L/5 and contact range is Long+Medium.

| Target | Precision Score (%) |
| --- | --- |
| T0866-D1 | 100.00 |
| T0912-D1 | 100.00 |
| T0944-D1 | 100.00 |
| T0896-D1 | 100.00 |
| T0894-D1 | 100.00 |
| T0896-D2 | 97.50 |
| T0947-D1 | 97.14 |
| T0942-D1 | 97.14 |
| T0892-D2 | 95.45 |
| T0882-D1 | 93.75 |
| T0948-D1 | 93.33 |
| T0946-D2 | 93.02 |
| T0918-D3 | 91.67 |
| T0909-D1 | 91.04 |
| T0907-D1 | 88.89 |
| T0913-D1 | 88.24 |
| T0868-D1 | 87.50 |
| T0892-D1 | 85.71 |
| T0911-D1 | 85.37 |
| T0942-D2 | 83.72 |
| T0912-D2 | 82.35 |
| T0898-D2 | 81.82 |
| T0887-D1 | 81.82 |
| T0894-D2 | 81.82 |
| T0918-D1 | 81.82 |
| T0886-D2 | 80.77 |
| T0899-D1 | 78.85 |
| T0864-D1 | 78.00 |
| T0901-D1 | 75.56 |
| T0907-D2 | 75.00 |
| T0895-D1 | 70.83 |
| T0898-D1 | 68.18 |
| T0872-D1 | 66.67 |
| T0876-D1 | 66.67 |
| T0890-D1 | 64.71 |
| T0915-D1 | 64.52 |
| T0900-D1 | 61.90 |
| T0904-D1 | 54.90 |
| T0878-D1 | 52.17 |
| T0905-D1 | 51.02 |

**Table S4.** List of targets with average precision value 50% or above when list size is Top10 and contact range is Long+Medium.

| **Target** | **Precision Score (%)** |
| --- | --- |
| T0894-D1 | 100.00 |
| T0946-D2 | 100.00 |
| T0892-D2 | 100.00 |
| T0942-D1 | 100.00 |
| T0866-D1 | 100.00 |
| T0944-D1 | 100.00 |
| T0942-D2 | 100.00 |
| T0912-D1 | 100.00 |
| T0896-D2 | 100.00 |
| T0896-D1 | 100.00 |
| T0909-D1 | 90.00 |
| T0912-D2 | 90.00 |
| T0864-D1 | 90.00 |
| T0868-D1 | 90.00 |
| T0947-D1 | 90.00 |
| T0948-D1 | 90.00 |
| T0882-D1 | 90.00 |
| T0886-D2 | 90.00 |
| T0892-D1 | 90.00 |
| T0918-D3 | 90.00 |
| T0899-D1 | 90.00 |
| T0918-D1 | 90.00 |
| T0901-D1 | 90.00 |
| T0913-D1 | 90.00 |
| T0907-D1 | 90.00 |
| T0894-D2 | 80.00 |
| T0898-D2 | 80.00 |
| T0911-D1 | 80.00 |
| T0907-D2 | 80.00 |
| T0907-D3 | 80.00 |
| T0890-D1 | 80.00 |
| T0878-D1 | 80.00 |
| T0915-D1 | 80.00 |
| T0895-D1 | 70.00 |
| T0918-D2 | 70.00 |
| T0898-D1 | 70.00 |
| T0941-D1 | 70.00 |
| T0904-D1 | 70.00 |
| T0872-D1 | 60.00 |
| T0888-D1 | 60.00 |
| T0876-D1 | 60.00 |
| T0887-D1 | 60.00 |
| T0870-D1 | 60.00 |
| T0900-D1 | 50.00 |
| T0874-D1 | 50.00 |

